# Single Cell Analysis of Peripheral TB-Associated Granulomatous Lymphadenitis

**DOI:** 10.1101/2024.05.28.596301

**Authors:** Philip J. Moos, Allison F. Carey, Jacklyn Joseph, Stephanie Kialo, Joe Norrie, Julie M. Moyarelce, Anthony Amof, Hans Nogua, Albebson L. Lim, Louis R. Barrows

## Abstract

We successfully employed a single cell RNA sequencing (scRNA-seq) approach to describe the cells and the communication networks characterizing granulomatous lymph nodes of TB patients. When mapping cells from individual patient samples, clustered based on their transcriptome similarities, we uniformly identify several cell types that known to characterize human and non-human primate granulomas. Whether high or low Mtb burden, we find the T cell cluster to be one of the most abundant. Many cells expressing T cell markers are clearly quantifiable within this CD3 expressing cluster. Other cell clusters that are uniformly detected, but that vary dramatically in abundance amongst the individual patient samples, are the B cell, plasma cell and macrophage/dendrocyte and NK cell clusters. When we combine all our scRNA-seq data from our current 23 patients (in order to add power to cell cluster identification in patient samples with fewer cells), we distinguish T, macrophage, dendrocyte and plasma cell subclusters, each with distinct signaling activities. The sizes of these subclusters also varies dramatically amongst the individual patients. In comparing FNA composition we noted trends in which T cell populations and macrophage/dendrocyte populations were negatively correlated with NK cell populations.

In addition, we also discovered that the scRNA-seq pipeline, designed for quantification of human cell mRNA, also detects Mtb RNA transcripts and associates them with their host cell’s transcriptome, thus identifying individual infected cells. We hypothesize that the number of detected bacterial transcript reads provides a measure of Mtb burden, as does the number of Mtb-infected cells. The number of infected cells also varies dramatically in abundance amongst the patient samples. CellChat analysis identified predominating signaling pathways amongst the cells comprising the various granulomas, including many interactions between stromal or endothelial cells and the other component cells, such as Collagen, FN1 and Laminin,. In addition, other more selective communications pathways, including MIF, MHC-1, MHC-2, APP, CD 22, CD45, and others, are identified as originating or being received by individual immune cell components.

**Author Summary:** The research conducted describes the cellular composition and communication networks within granulomatous lymph nodes of tuberculosis (TB) patients, employing a single-cell RNA sequencing (scRNA-seq) approach. By analyzing individual patient samples and clustering cells based on their transcriptome similarities, the study reveals several consistent cell types described to be present in both human and non-human primate granulomas. Notably, T cell clusters emerge as abundant in most samples. Additionally, variations in the abundance of B cells, plasma cells, macrophages/dendrocytes, and NK cells among patient samples are observed. Pooling scRNA-seq data from 23 patients enabled the identification of T, macrophage, dendrocyte, and plasma cell subclusters, each displaying distinct signaling activities. Moreover, the study uncovers a surprising capability of the scRNA-seq pipeline to detect Mtb RNA transcripts within host cells, providing insights into individual infected cells and Mtb burden. CellChat analysis unveils predominant signaling pathways within granulomas, highlighting interactions between stromal/endothelial cells and other immune cell components. Moreover, selective communication pathways involving molecules such as Collagen, FN1, Laminin, CD99, MIF, MHC-1, APP and CD45 are identified, shedding light on the intricate interplay within granulomatous lymph nodes during TB infection.

## Introduction

Lymphadenopathy refers to the swelling of lymph nodes that can be secondary to bacterial, viral, or fungal infections, autoimmune disease, and malignancy [1, 2]. Once malignancy is ruled out, the association of the lymphadenopathy with a medical history and symptoms consistent with tuberculosis (TB) is a key diagnostic consideration. Granulomatous lymphadenopathy caused by *Mycobacterium tuberculosis* (Mtb) is a signature feature of tuberculosis. Such granulomas are composed of immune and non-immune cells that are thought to mount an orchestrated pro- and anti-inflammatory responses to the bacterium [3-12]. Long deemed protective to the host, the granuloma is now understood to benefit the bacterium in some ways and represent a site of critical host-pathogen interactions that drive infection outcomes [12]. We present here the first single cell transcriptomic analysis of human peripheral tuberculosis–associated granulomatous lymphadenopathies. This analysis highlights the contrast in information gained by single cell analysis, as compared to established histological approaches (9, 10).

TB remains a major global health emergency, causing over 1.5 million deaths in 2022 [13-16]. In low-resource settings, the diagnosis of TB is often based on history and symptomology, supported by acid-fast Ziehl–Neelsen staining microscopy of sputum or biopsy smear, and GeneXpert molecular testing when available [17,18]. It is estimated that 50% or less of all active TB cases are identified by microscopy in such resource limited settings [19-21]. At the TB clinic of Port Moresby General Hospital, in Papua New Guinea, it is standard protocol to take fine needle aspirates of peripheral lymph nodes greater than 0.8 cm in diameter in presenting TB patients, in order to obtain samples for microscopy and GeneXpert analysis, a molecular diagnostic.

We obtained FNA samples from such TB patients (S-TABLE 1) and report here the single-cell RNA sequencing (scRNA-seq) characterization of the cell types detected and analysis of their communication networks. When mapping cells from individual patient samples, clustered based on their transcriptome similarities, we uniformly identify several cell types that are known to comprise granulomatous lymphadenopathies in humans and non-human primates (NHPs) [22-24]. We find the T cell cluster to be one of the most consistent and abundant. Other cell clusters that are uniformly detected, but which vary dramatically in abundance amongst the individual patient samples, were the B cell, plasma cell, macrophage/dendritic cell and Natural Killer (NK) cell clusters. When we combined all the scRNA-seq data from our current 23 patients (in order to add power to cell cluster identification in patient samples with fewer cells or of poorer quality), we distinguish B, T, macrophage, dendritic cell and plasma cell subclusters, each with distinct signaling activities. The sizes of these subclusters also varies dramatically amongst the individual patients suggesting that the predominance of their respective immune signaling pathways of bacterial control may vary proportionately. In analyzing the cellular composition of our highest quality samples, we defined some statistically significant trends in the FNA cellular composition, the most significant being that as the percentage of Natural Killer cells increases, T cell numbers decrease. Upon analysis of T cell markers, it was noted that CD8A and IL7R positive T cells decrease with the NK cell increase. A less significant trend that was identified suggests an increase in macrophage and dendrocyte cells might increase as NK cells increase and T cell percentages decrease.

In a surprising result, we also discovered that the scRNA-seq pipeline [25], designed for quantification of human mRNA, also detects Mtb RNA transcripts and associates them with their infected-host cell’s transcriptome, thus identifying individual infected cells. The numbers of Mtb reads and infected cells also vary dramatically in abundance amongst the patient samples. We postulate that the number of detected bacterial transcript reads provides a measure of Mtb burden, as does the number of Mtb-infected cells. It is possible that the number of Mtb transcript reads, or the number of infected host cells, can distinguish high versus low Mtb burden granulomas. In comparing differentially expressed features by heat map analysis, it was noted that MALAT1 [26, 27], a long, non-coding RNA is significantly down-regulated in TB-infected cells, among other changes. Thus, this first of-its kind set of human TB FNA samples has allowed us to identify cell-cell communication ligands and receptors likely to play important roles in the functioning of the presumed human peripheral lymph node granuloma. Further, it has identified trends in granuloma composition that may correlate with their relative inflammatory status. This initial single cell analysis of 23 human FNAs is consistent with literature suggesting a great deal of variation in cellular composition and bacterial load in FNA samples from TB patients with peripheral lymphadenopathy.

## Results

### Determination of FNA cellular composition

In 2016 we began a collaboration with Dr. Evelyn Lavu (decd.) and colleagues at the Port Moresby General Hospital (POMGH) Tuberculosis Clinic, Papua New Guinea, to obtain an MRC-approved (IRB) protocol that allowed us to develop our protocol for scRNA-seq analysis of human FNA samples from peripheral nodal granulomas of tuberculosis patients. At the POMGH, TB patients referred to the clinic are assessed based on presentation, symptoms and history. Fine needle aspiration is the established protocol to assist diagnosis of patients exhibiting accessible lymph node granulomas exceeding 0.8 cm diameter. For patients agreeing to this assessment, three fine needle aspirate passes of the granuloma are harvested for acid-fast microscopy and for GeneXpert analysis [17,18]. Our protocol provided us one of these aspirate samples, which was preserved in methanol for sterilization, transportation and subsequent re-hydration. After several rounds of methods optimization, we have accomplished scRNA-seq analysis of FNA samples from 23 individual patients.

In 2019 we began to obtain de-identified FNA samples from permission-granting patients. Only one sample from 2019 yielded a library of sufficient quality for sequencing (FNA 0). From 2022, using improved protocols we obtained libraries from six FNA samples, however most of these (FNAs 1, 2, 3, 4 and 8) (Fig.1) had high numbers of cells that were essentially unidentifiable when analyzed individually because of the lack of discriminating transcripts. With further improvement of our protocol (see Methods), we had much better success with FNAs collected in 2023, obtaining high-quality libraries and sufficient numbers of cells from most of the FNAs 10 through 28.

**Fig. 1.**
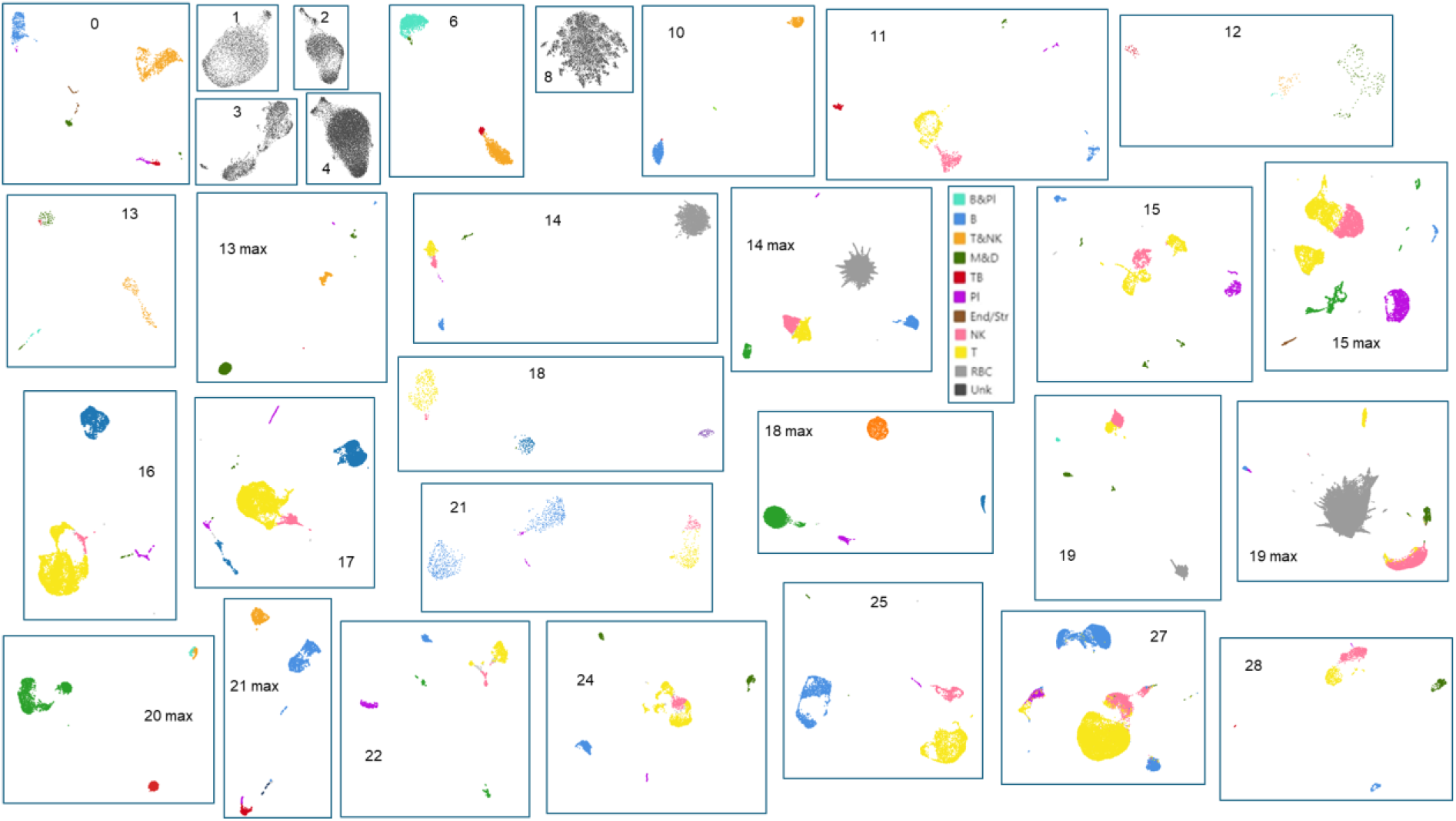
Individual FNA UMAPs and Cell Cluster Identifications. CellRanger indexed BAM files as cloupe.cloupe files were visualized using the 10XGenomics Loupe Browser 7.0.1. Reads were mapped to human (hg38), H37Rv Mtb (NC_00096.3:1) and HIV-1 (NC_001802.1) genomes and dimension reduction accomplished using the Loupe Browser designating 30 PCAs and clustering to the nearest 10 cells. FNA number show in each panel. Several FNA libraries (FNA 1, 2, 3, 4 and 8) were too poor to cluster when analyzed individually (cells designated as unknown), although they all contained Mtb positive cells and other identifiable cellular components when analyzed in the larger combined data set (Fig. 6). Cell types constituting the recognized clusters for each FNA were identified by the differential expression of selective marker transcripts (Table 1). Some FNAs with lower quality or fewer numbers of cells were run through the entire 10XGenomics pipeline a second time (designated max) starting with the maximum possible volume of cells for analysis. For FNA 10, the RNA had degraded and yielded a poor quality max library during the re-run, and is not included here. For FNA 20, the initial library was too small to include as well. Significant erythrocyte contamination is apparent in FNA 14 and 14 max, and FNA 19 and 19 max. For the 10 UMAPS of highest quality, T and NK cells could be enumerated in separate clusters. The other FNA samples, with less resolution, showed intermingled T and NK or intermingled B and plasma cells (clusters T&NK and B&Pl, respectively), which were enumerated as such to calculate the FNA cellular compositions shown in figure 2.

**Table 1.** Selectively expressed cell marker genes used for cluster-cell type identification, with other tools.

| Selectively Expressed Cell Marker Transcripts |  |  |  |
| --- | --- | --- | --- |
| Name | Cell Type | Name | Cell Type |
| CD4 | T helper cells | LILRA4 | Dendrocytes |
| CD3D | T cells | LAMP5 | Dendrocytes |
| CD8A | T cytotoxic cells | LRRC26 | Dendrocytes |
| TIGIT | T cells | FDCSP | Germinal dendrocytic cells |
| FOXP3 | T repressor cells | ACSL1 | Foam cell |
| IL7RA | T cells | LTA4H | Caseous |
| GZMA | T cytotoxic cells, NK cells | RNASE1 | Macrophages, endothelial |
| TBX21 | Th1 cells, NK cells | COL3A1 | Endothelial |
| NKG7 | NK cells | COL4A1 | Endothelial |
| CX3CR1 | NK cells | CD31 | Endothelial, monocytes, adipocytes |
| FUT4 | Neutrophils | EMP1 | Endothelial, keratinocytes, adipocytes |
| CD14 | Macrophages | EPAS1 | Endothelial, smooth muscle, adipocytes |
| CD163 | Macrophages | CD34 | Endothelial, smooth muscle, adipocytes, glandular cells |
| APOC1 | Macrophages | CD40 | B cells |
| CSF1R | Macrophages | CD79A | B cells |
| CD68 | Macrophages | CD24 | B cells |
| CPVL | Macrophages | MS4A1 | B cells |
| LYZ | Macrophages | CD19 | B cells, plasma cells |
| C1QB | Macrophages | IGHG1 | Plasma cells |
| LILRA1 | Macrophages | JCHAIN | Plasma cells |
| LILRA5 | Macrophages | HBA1 | RBC |
| CLEC4C | Dendrocytes | HBB | RBC |
| SCT | Dendrocytes | HBG2 | RBC |

After rehydration and processing through the 10X Genomics pipeline for scRNA-seq and Illumina sequencing, the transcriptomes were assembled, annotated and data visualized using Cell Ranger and Seurat v4/tSne or UMAP analyses and dimension reduction tools (see Materials and Methods) to yield the 10XGenomics Loupe Browser 7.0.1 UMAPs shown (Fig. 1). Because of the low cell numbers detected in the initial analyses, some of the FNAs were re-run through the entire 10X Genomics pipeline at the maximum cell volume (designated max) when both the initial and the -max re-runs yielded libraries of sufficient quality, the data were combined. The differential expression of selective cellular transcripts (Table 1), obtained from the literature and 10X Genomics Loupe Browser 7.0.1 heat map analyses (selectivity vetted using the Human Protein Atlas website [28]) was used to identify and quantify the cell types detected in the FNA samples in order to determine their cellular make up. CellMarker 2.0 [29] and SingleR 1.12.1 [30] analysis of the combined data set (below), were also used to help identify the cell types as shown (Fig. 1 and 2.). While each patient sample was different, with FNA volumes as low as ∼100 µL, the numbers of cells recovered after preservation in methanol ranged narrowly, from approximately 7X10^6^/mL to 6X10^7^/mL. This provided an excess of cells for scRNA-seq analysis and allowed re-running specified samples.

**Fig. 2.**
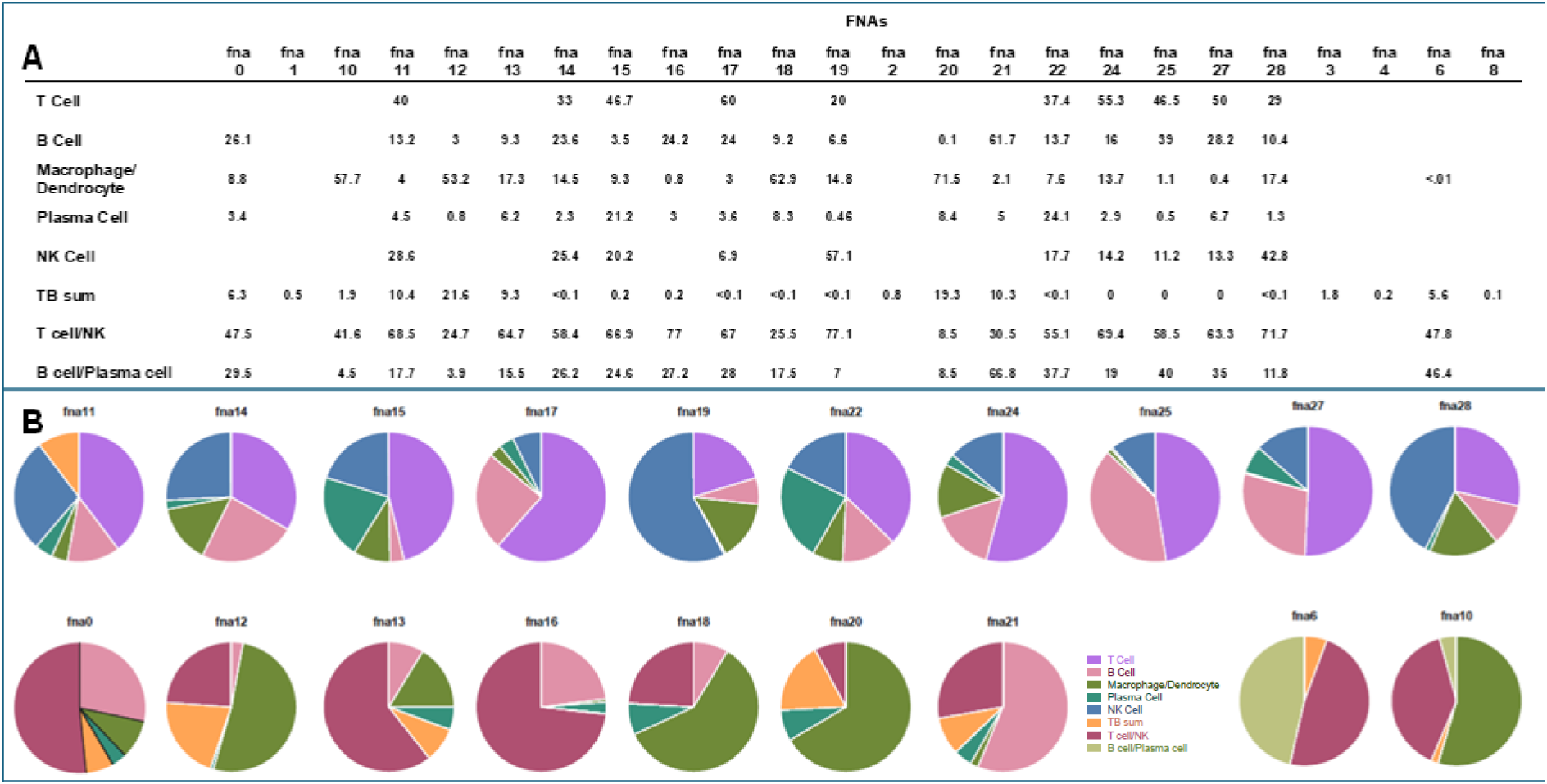
Cell Composition of Individual FNA Clusters. **Panel A)** A matrix of identified cell cluster percentages making up the individual FNAs as resolved by the Loupe browser and identified by differential expression of marker transcripts (from Fig. 1). Raw cell count totals are presented in (S-1). **Panel B)** Pie charts of FNA composition. Upper panel of pie charts shows FNAs for which NK cell clusters could be quantified separately from T cells (FNA 11, 14, 15, 17, 19, 22, 24, 25, 27 and 28). Cluster identities were later confirmed by algorithm driven cell clustering in the combined data set. Lower panel of pie charts shows the FNAs with intermingled T and NK cell clusters (FNA 0, 12, 13, 16, 18, 20 and 21), and those also with intermingled B and plasma cell clusters (FNA 6 and 10).

When analyzed individually using the Loupe browser or Seurat dimension reduction tools, most of the FNA samples yielded UMAPs with a limited numbers of clusters that included most of the component cells, usually easily attributable to given cell types based on marker transcripts, (Fig. 1). Most commonly identifiable was a T cell cluster (CD3D, IL7R, CD8A, etc., positive). The T cell cluster was often intermingled with NKG7 (natural killer) cells, depending on the number of cells and the quality of the library (Fig. 2). When this was the case the cluster was designated T&NK, rather than clearly definable as individual T and NK clusters. Other clusters routinely detected included a B cell cluster (MS4A1, CD40, CD79A, etc., positive cells), a plasma cell cluster (JCHAIN and IGHG1 positive), a macrophage/dendrocyte cluster (CD14, CD68, CD163, CPVL, LYZ, CEC4C, LILRA4, etc., positive), and a TB positive cluster (Mtb rrs and/or rrl transcript positive). It was clear that while the T cell cluster was uniformly detected as one of the most abundant harvested from every patient, the other cell type clusters varied dramatically (Fig. 2). As mentioned, in some FNAs with poor quality libraries, or with fewer cells, individual UMAPs could not distinguish meaningfully amongst the detected T and NK, or B and plasma cells. Cells expressing many B or T cell subtype, or macrophage/dendrocyte differential subtype markers were routinely intermingled into the single the B, T or macrophage/dendrocyte Loupe Browser UMAP clusters. Cell numbers varied from a low of 231 for FNA 12 to a maximum of 26, 804 for FNA 27 (S-1). Only infrequently did we detect clusters of cells that are identifiable as something other than T, B, plasma, macrophage/monocyte or dendrocyte types (e.g., cells expressing “selective markers” for putative erythroblasts or endothelial cells). We did not detect neutrophils that clustered separately, probably because they are known to be relatively transcriptionally inactive and not harvested in significant number by fine needle aspiration. SingleR analysis suggests they cluster with other phagocytes (S-2).

We first sought to define the cellular composition of the 23 FNA samples and to look for any trends in that data. Multiple approaches were taken to quantify cell clusters generated in the Loupe Browser UMAPS. Initially, barcodes of cells displaying a selective marker were quantified and summed to 100 percent for a given UMAP. However, with the relatively low coverage we found in some of our samples the vast majority of cells went uncounted.

Furthermore, it was uncertain whether a given marker for B cells would yield barcode numbers as efficiently as a given marker for T cells. As an alternative, the barcodes for all the cells comprising the identified cell type clusters for each sample UMAP (Fig. 1) were determined individually, using K means differentiation and the Loupe Browser lasso tool to quantify the cells comprising a given cell type cluster. These cells were then summed to determine the percentage contribution of each cluster to the cell total in a given UMAP (Fig. 2). It was these percentages that were used to generate the pie charts shown in Figure 2.

Initial analyses tracking cellular composition versus increasing Mtb-infected cells did not reveal any significant trends. Neither did other analyses tracking FNA composition versus increasing numbers of any of the other comprising cell types when the T and NK cluster percentages were combined (e.g., samples with T&NK clusters in Fig. 2). However, when we focused on the 10 higher quality FNAs in which a T cell cluster could be distinguished from the NK cell cluster, a trend was identified in which increasing NK cell percentage was inversely related to the number of T cells (Fig. 3). A second, weaker trend was identified, in which there is a suggestion that the macrophage/dendrocyte cell percentages increases proportionately with NK cell number in the sample set. To further assess the inverse relationship between T cells and NK cells, as they comprise these FNA samples, we tracked our selective T cell marker barcodes versus increasing NK cell barcode numbers. It was found that, as the percentage of NKG7 or GZMA positive cells increased, the percentage of CD8A and IL7R positive cells decreased, suggesting a decrease in Th1-proinflammatory T cell presence as NK cell number increases (or vice versa) (Fig. 4).

**Fig. 3.**
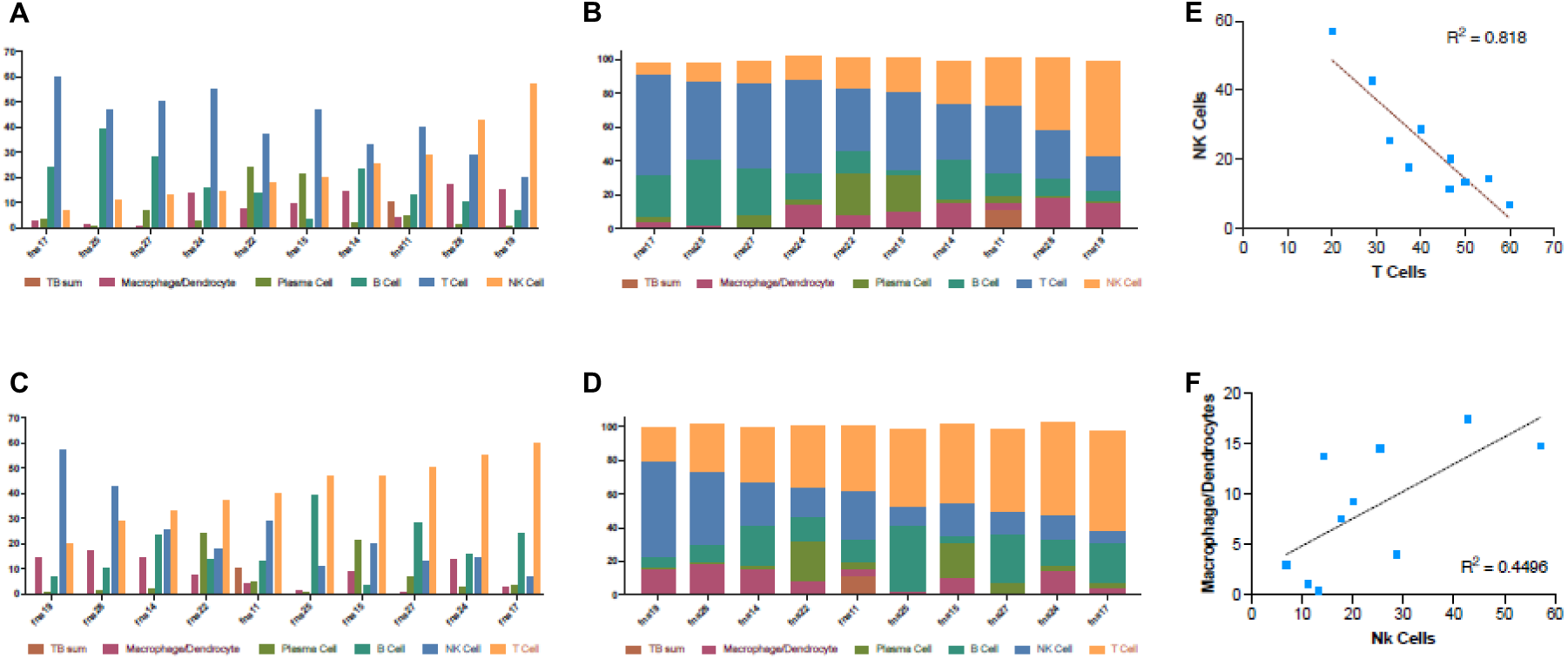
Trends in Cellular Composition of FNA samples. The percentage of cell types making up given FNAs (from Fig. 2) were plotted as histograms or stacked bar charts to identify obvious trends (**Panels A and B** show others cell percentages versus increasing NK cells; and **Panels C and D** show others versus increasing T cells). Trends suggested in panels A-D were then evaluated by fitting trend lines to scatter plots of the data (**Panels E and F**). Significant trends were identified suggesting an inverse relationship between the numbers of NK cells and T cells, and a positive relationship between numbers of NK cells and macrophage and dendrocyte cluster cells.

**Fig. 4.**
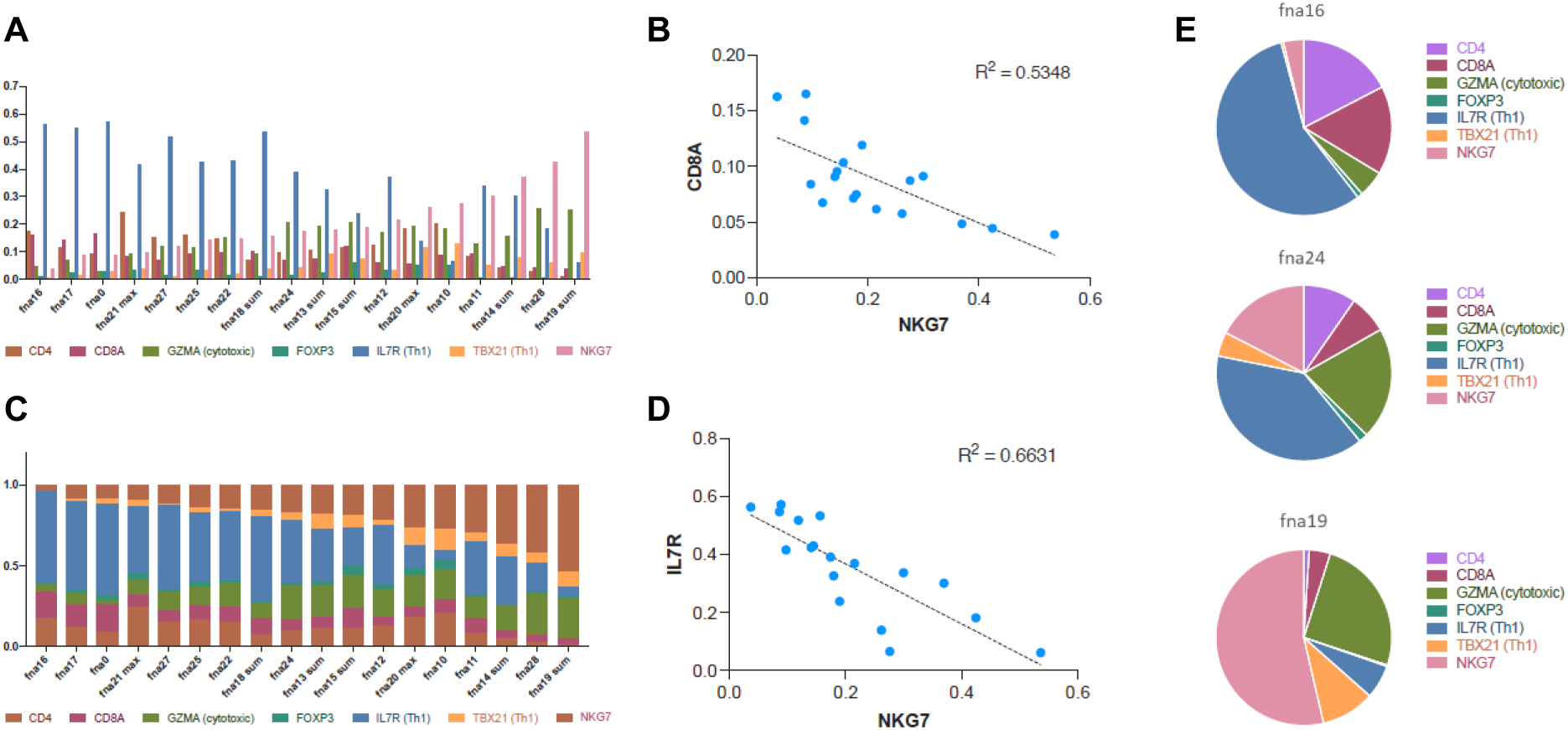
Trends in T cell Composition versus NK cell Population. Histogram and stacked bar charts **(Panels A and B)** were used to visualize trends between various T cell markers and increasing numbers of NK cells (which correlate with decrease T cell populations overall, Fig 3). Significant trends (**Panels C and D**) suggest that T cells with CD8A and IL7R markers are the T cell populations that decrease most significantly as the percentage of NK cells in the FNA increases. Pie charts of FNA 16, 24 and 19 are presented (**Panel E**) to illustrate the trends.

### Cellular Communications Indicated by the Combined Data Set

To increase our power, and to attain greater precision in cell type identification, we decided to combine all the patient data into one UMAP in order to gain sufficient power for cell type identification in the samples of poorer quality or lower numbers of cells (Fig. 5) and (S-3). This increased number would also empower meaningful description of cell-cell communication networks of cells characterizing the combined FNA data set. While requiring greatly increased computing power, combining the over 202,000 cell transcriptomes from our 23 patient FNAs did provide increased confidence in the ability to visualize the distribution of cell types amongst our various patient samples (Fig. 6). In the poorer quality FNAs, with large numbers of un-identifiable cells, there were still minor clusters of cells with sufficient similarity to the combined clusters for confident cell type identification. Attempts to assign identities to cells making up Cluster 2 using SingleR or other tools yielded only varying embryonic and neuronal cell type assignments. Significantly, several subsets of the major cell types (B, T, plasma and macrophage/dendrocyte) were resolved that evidenced differential cellular communication networks (Fig. 8). When the individual FNA UMAPS were extracted from the combined data set, it was again apparent that many FNAs exhibited significant differences in their cellular composition (Fig. 6), in agreement with the individual analyses shown in figures 1 and 2.

**Fig. 5.**
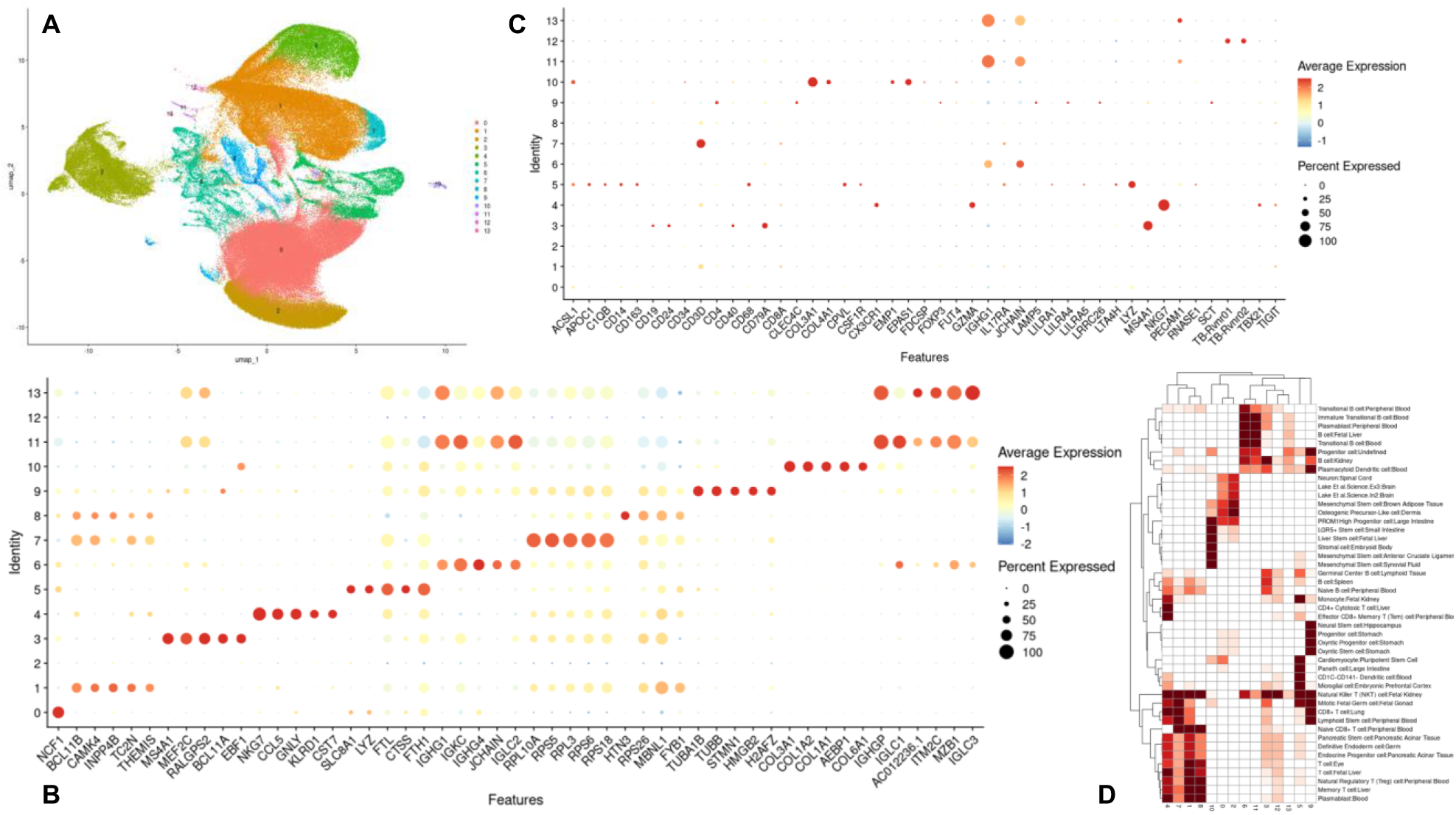
Combined Data Set Cluster Identification. Combination of all 202,000 cells from our combined 23 FNAs (initial and max reruns combined) in SEURAT analysis designating a resolution of 0.1, yielded the UMAP shown (**Panel A**). Cluster 2, consisting of unidentifiable cells largely contributed by FNAs of poor quality (see Fig. 6), and was removed from the data set when plotted against the expressed features. The top 5 most differentially expressed genes (**Panel B**) and expression of our selective marker genes (**Panel C**) were used in combination with SingleR (S-3), CellMarker_Augmented_2021 (**Panel D**) and other tools, to identify cell types making up the 14 clusters identified in the combined data set at this resolution. Higher resolution was found to be overly divisive and not informative. Identification of the clusters shows that most of the general cell types identified by Loupe browser clustering (Fig. 1) were divided into subtypes (all except for NK cells). Clusters are identified as: 0=pre-macrophage; 1=T(1); 3=B; 4=NK; 5=macrophage; 6=plasma(1); 7=T(2); 8=T(3); 9=dendrocyte; 10=stromal/endothelial; 11=plasma(2); 12=TB infected; 3=plasma(3). CellChat analysis shows these sub-clusters express distinct cell-cell communication ligands or receptors (Fig. 8).

**FIG. 6.**
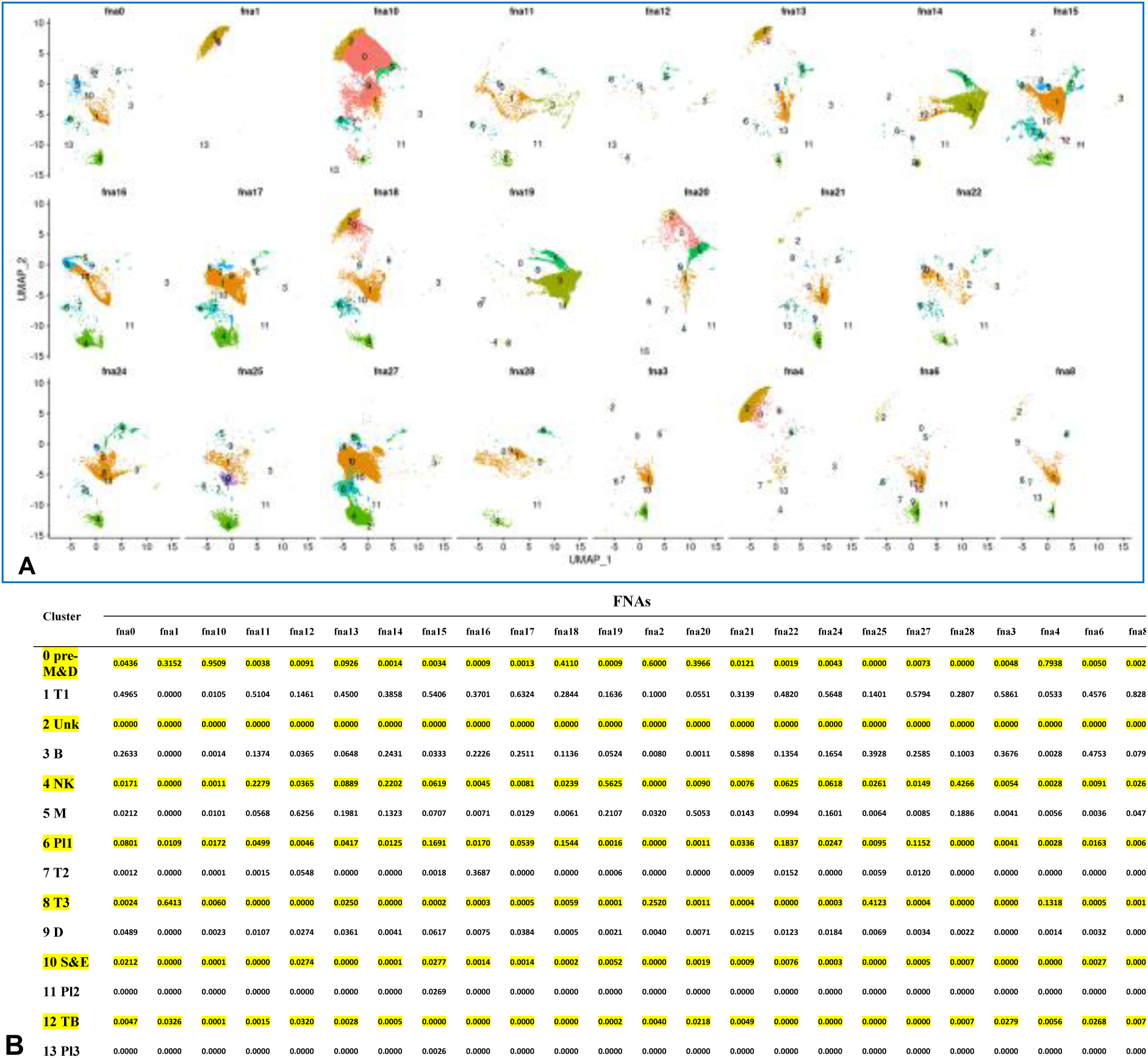
Composition of Individual FNAs Determined from the Combined Data Set. (**Panel A**) The individual FNA UMAPS extracted from the Combined UMAP (from Fig. 5) show that the strategy to combine all our patient samples in order to increase the precision with which of cell types, or sub-types, could identified was successful. In fact, identifiable cell were obtained from all the patients FNAs, even those of poor quality that had large percentages of unidentifiable cells. Here, while FNAs 1, 2, 3, 4 and 8 still have large numbers of unidentifiable cells (Custer 2), as did FNA 10, 13, 18 and 20. Nevertheless, these poorer quality samples provided significant, though varying, numbers of identifiable cells to the combined set, which empowered CellChat analysis (Below). A matrix of the percentages of each of the identified cell clusters in shown in **Panel B**. Some of the minor cell cluster (e.g., Cluster 11 and 13-plasma cells) are often identified in only one or two of the FNAs. Raw numbers of cells making up the extracted FNA UMAPS shown in figure S-3.

**Fig 7.**
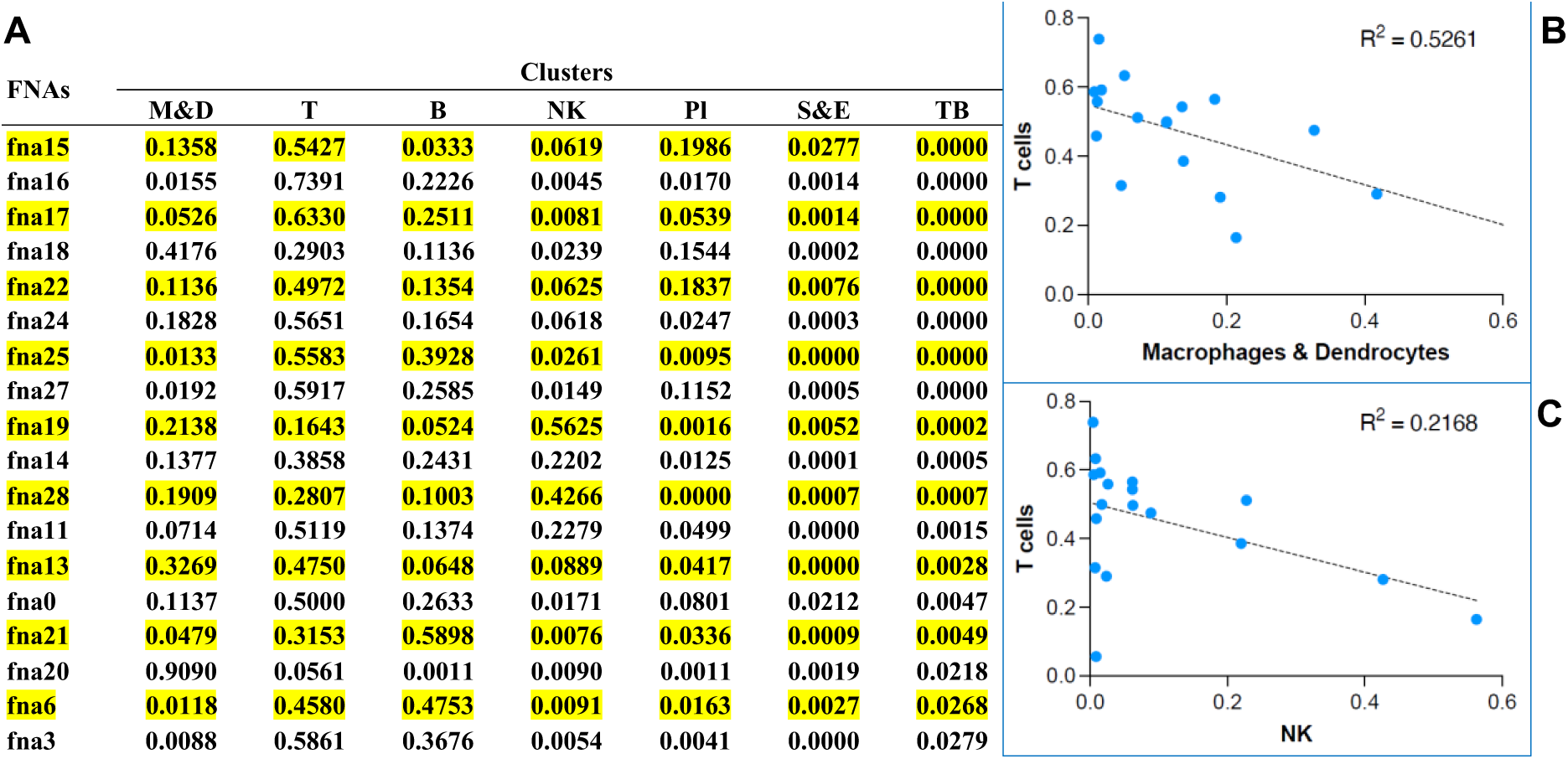
Trends in Cellular Composition of FNA Samples Using Combined Data Set Clustering and Cell Identifications. T, macrophage and dendrocyte, and plasma cell cluster percentages from Figure 6 were summed (**Panel A**) to determine percentages of “parental” cell types making up the given FNAs. The cell type percentages from FNAs containing more than 1,000 cells were then plotted to visualize any trends in FNA cell type composition (**Panels B and C**). An obvious trend suggested by the whole combined data set was a decrease in macrophage/dendrocyte cell numbers as T cell numbers increase (**Panel B**), reflecting a trend identified from the Loupe Browser UMAP data (Fig. 3). The previously identified correlation of an inverse relationship between NK and T cell cluster sizes was also apparent, although with less certainty when plotted using this combined data set (**Panel C**).

**Fig. 8.**
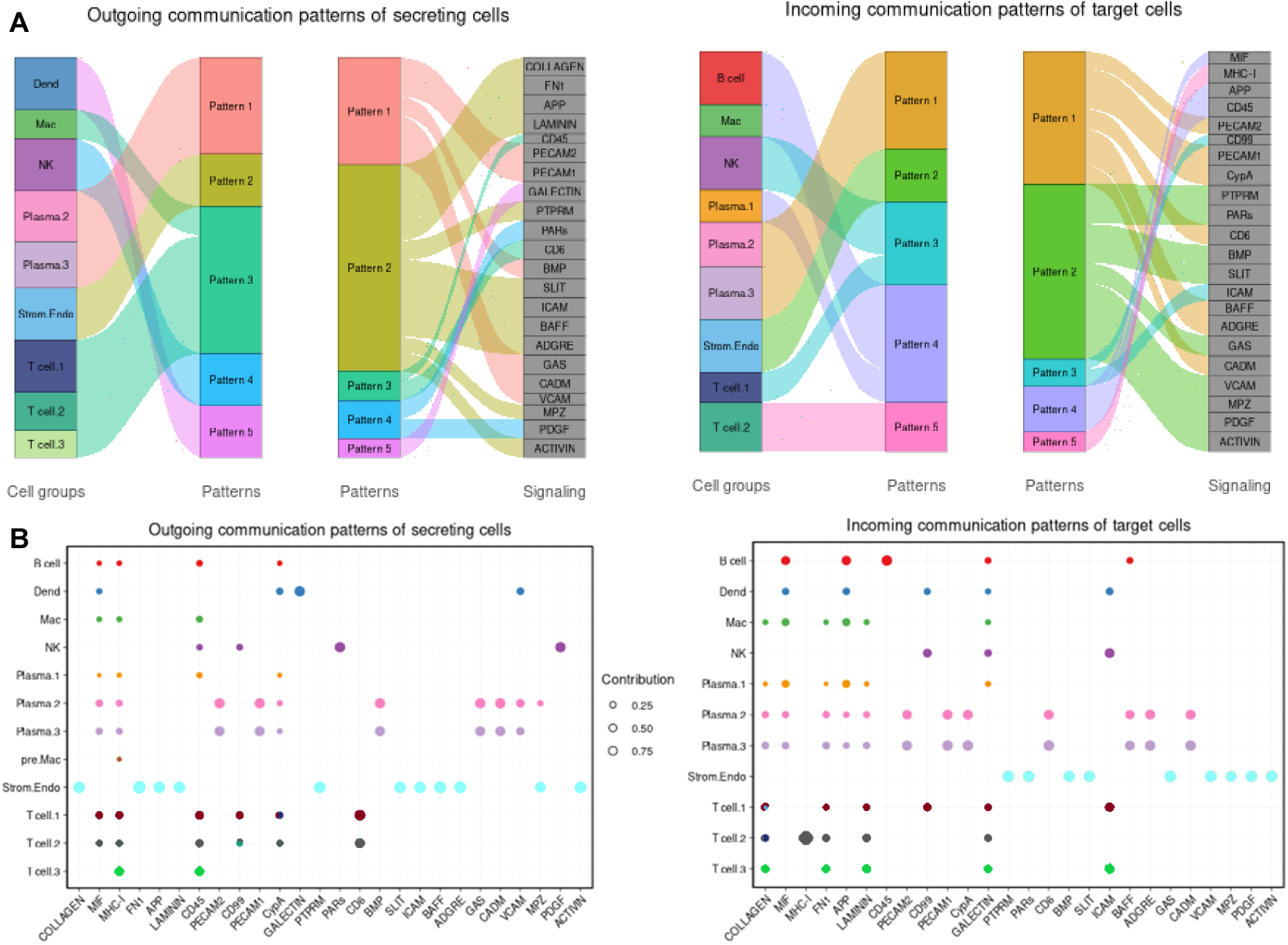
CellChat Signaling Interactions for Combined FNA DATA Set. The patterns of signaling interactions between the combined data set clusters (Fig. 5) and their relative strengths are shown in **Panel A**. **Panel B** shows “Outgoing” and “Incoming” communication ligands and receptors for the clusters identified from the combined data set. Cluster 10, (stromal/endothelial) cells are identified as unique senders and receivers for many communication patterns. Dendritic cells appear to be unique senders in Galectin signaling. NK cells are identified as unique senders in PARs and PDGF signaling. The T cell 2 cluster is identified as the unique recipient of MHC-1 signaling, and B cells are identified as unique recipients for CD45 signaling. Most cell clusters are identified as sharing multiple signaling patterns with other cell types in the FNAs. The predominating signaling pathways identified varied with CellChat version 1 (S-6 shows CellChat version 1.1 analysis of our original combined data set, before default minimization of high Hb gene and MT gene cells in the combined set shown here).

As shown in figure 6, the composition of individual FNAs could be extracted from the Combined Data Set. The clusters are ranked by cell number, Cluster 0 being the most populous, Cluster 14 being the least populous. As an independent, fully algorithm-driven dimension reduction-clustering result, we asked if the ratios of the major T, B, macrophage/dendrocyte, NK, plasma and TB cell cluster percentages agreed with those determined by the individual Loupe analyses shown in figure 1. When the respective cell type sub-clusters from the combined data were summed and the percentages of the total determined, there was close agreement of the FNA compositions extracted from the combined data set with the FNA compositions determined by the individual Loupe analyses (Fig. 2). This allowed us to ask if the trends in FNA composition, identified from the Loupe Browser analyses, were repeated in this larger data set, which included cell contributions from the FNAs of poorer quality and relied on the power of the increased N for cell type identification. Basically, we questioned if the inclusion of increased numbers of poorer quality cells, identified because of the increased power of the analysis, confirmed or undermined the trends identified using numbers from the individual Loupe analyses. This analysis of the combined data set did confirm trends identified in figure 3, with caveats. A significant trend showing decreased macrophage/dendrocyte cells versus increasing T cell percentages was identified in the complete combined data set (Fig. 7). In contrast, the correlation of the inverse relationship between NK and T cells was less significant. However, if the FNAs of poorer quality or number were excluded (i.e., those with a large cluster 2 or low numbers of cells), the previously identified inverse relationship between NK and T cell numbers and proportional relationship between NK and macrophage and dendrocyte cells confirmed (Fig. 7). We felt the exclusion of these poorer quality FNAs was justified, because it was found that the FNAs, 1, 2, 4, 8 and 12, extracted from the combined data set, had only a total of 92, 27, 732, 901 and 219 identifiable cells each (respectively), thus contributing to greater variation amongst the clusters detected in each. In addition, in the combined data set all the FNAs conducted with a “max” repeat were combined regardless of quality. This included the “max” repeats for FNA 10max (combined with FNA 10) and the initial FNA 20 (combined with FNA 20 max) samples, which by individual Loupe Browser assessment had failed and did not yield quality libraries. Thus, we feel that the FNA composition trends shown in Figs 3, 4 and 7 appear to repeat and to be significant within this limited data set.

The CellChat tool was used to predict cell communication ligands and receptors expressed in the combined data set clusters. “CellChat quantifies the signaling communication probability between two cell groups using a simplified mass action-based model, which incorporates the core interaction between ligands and receptors with multi-subunit structure along with modulation by cofactors” [31]. Cell-Cell communications of the various B, T, macrophage/dendrocyte and plasma cell subclusters suggests that Seurat UMAP clustering of the combined data set did in fact distinguish amongst cell type subsets based to some extent on their signaling status. When checked, it was found that these same “Sender and Receiver” ligands and receptors transcript markers, designated as selective for a particular cell type subset from the combined data, mapped to the appropriate respective cell type cluster in the individual Loupe Browser UMAPS. Thus, a cluster 0 (pre-macrophage and dendrocyte cells) cell-specific ligand transcript from the combined data set, maps to the single macrophage/dendrocyte cell cluster identified in the Loupe Browser UMAP of individual FNA analyses, and so on. This again supported the consistency between our two approaches to the FNA analysis, individual Loupe Browser versus the combined data set analysis using the Seurat UMAP tools.

The current Combined Data Set used for cell signaling analysis in Fig. 8 used default removal of hemoglobin or mitochondrial expressing cells in the Cell Ranger/Seurat assembly and the CellChat V2. It was seen that the number of CellChat cell communication interactions identified were different from those obtained with our Original Combined Data Set analyzed using CellChat v1 (S-4, 5, 6). The principle difference between the two UMAPS in terms of cell cluster identification, was the identification of a new individual endothelial/stromal cell cluster (cluster 10) in the noHB data set (Fig. 5). Analysis of Excel files of cluster specific UMAP features showed that these COL3 and COL4 expressing cells (endothelial/fibroblast/smooth muscle) were mostly incorporated into the pre-macrophage and dendrocyte cluster (Cluster 0) of the original data set (S-2). Cell-cell communications identified by CellChat are clearly influenced by the QC defaults set during analysis as well as CellChat version.

CellChat analysis of the combined data set ranked Collagen, MIF, MHC-I, FN1, APP, and Laminin as the top 6 signaling interactions. Collagen signaling monitors the extracellular matrix while helping define cell shape and behavior. MIF (macrophage migration inhibitory factor) involves cell-cell contact between many cell types, but predominately, T and plasma cells as senders, and the pre-macrophage/dendrocyte cluster (Cluster 0) and B cell cluster (Cluster 3) as receivers (Fig. 8). The MHC-I signaling, as defined by CellChat, identified multiple plasma, pre-macrophage/dendrocyte and T cluster cells as interacting with a specific T cell 2. FN1and Laminin signaling, like Collagen signaling, emphasize the overarching role the stromal endothelial components of the FNA in communicating with the other component cells of the granuloma. Signaling APP signaling is predominately from pre-macrophage and dendrocyte cluster 0 cells to cluster B cells and themselves.

CellChat v1analysis of the Original Combined data set ranked MIF, MHC-II, MHC-I, APP, CD22 and CD45 signaling as among the top interactions. MHC-II signaling identified principally cluster B cells and macrophage cells and pre-macrophage cell cluster as senders, with pre-macrophage cluster cells as the principal receivers. In the original combined data set, CD22 signaling was shown to be predominately from the B cell cluster to multiple T, B and macrophage/dendrocyte receivers, while CD45 signaling was the reverse, from multiple cell types to the B cell cluster, etc.

It was interesting to note that some of the smallest clusters of cells (stromal/endothelial, plasma 2 and 3, T 2 and 3 and B 2) appeared to be driving the identified cell-cell communications. We postulate that these small clusters appear to be separated out in the Seurat analysis specifically because they are very active in one or another signaling system. An example would be the small plasma cell 2 and 3, which are almost exclusively found in FNA 15. The differences in the cellular composition of the given FNAs likely reflects the different predominating signaling networks in the respective granulomas.

### Status of Mtb-infected cells

We hypothesize that quantification of Mtb reads (Fig. 8), or of infected cells (e.g., Fig. 2; infected cells can contain more than 1 detected Mtb .sam read), will correlate with Mtb burden. The detection of bacterial transcript reads appears to be a stochastic process, with most of the detected sequences being from either the rrs or rrl small and large ribosomal subunit genes (Rvnr01 and Rvnr02, respectively) (Fig. 8). We hypothesize this is might be due to the high level of ribosomal rRNA transcripts expressed in the Mtb cells. However, it is possible that other features such as secondary structure and nucleotide modification could contribute to the spurious priming of these transcripts in the 10X Genomics pipeline, or perhaps even spurious priming with the UMI-leader sequence. The 10X Genomics pipeline we used relies on poly-T primers to initiate reverse transcription. Clearly, some patient FNAs have relatively high numbers of Mtb reads (thousands) detected, while other patient samples with only a few or none (Fig. 8). We had anticipated that trends of cellular composition in our FNA samples might correlate with increasing or decreasing numbers of Mtb infected cells in those samples. However, as reported above, we did not observe such trends in this limited data set when analyzed either using the individual FNA Loupe Browser or combined data set FNA compositions. Furthermore, when using CellChat to assess communications between cell type clusters, the TB-infected cluster (cluster 13) appeared to be virtually silent.

When looking for selective transcript features of Mtb-infected cells in an *in vitro* THP-1-GFP-H37Ra co-culture experiment (unpublished data), we noted that the long non-coding RNA, MALAT1, was routinely down-regulated in infected THP-1 cells. NEAT1 was found to be similarly down regulated, although less reliably in repeat experiments. Because of the possible immunomodulatory role of these long non-coding RNAs in viral infections [26, 27, 32] we asked if MALAT1 was downregulated in the TB-infected clusters of our patient FNAs. When assessing the 7 FNAs with highest overall numbers of Mtb-infected cells, we observed that in five of them (FNA 6, 11, 12, 13 and 21), MALAT1 was the most down-regulated transcript. In the other two FNAs assessed (0 and 20), MALAT1 was also significantly down regulated. The Expression of MALAT1 was then assessed in Mtb positive cells versus the other clusters identified in the Combined Data set (Fig. 10). This analysis confirmed down regulation of MALAT 1 in the Mtb infected cells across the complete data set. While many other transcriptional features vary amongst the FNA clusters, it is interesting that MALAT1 appears to stand out, perhaps suggesting a role in the bacterium-macrophage amour-haine relationship.

**Fig. 9.**
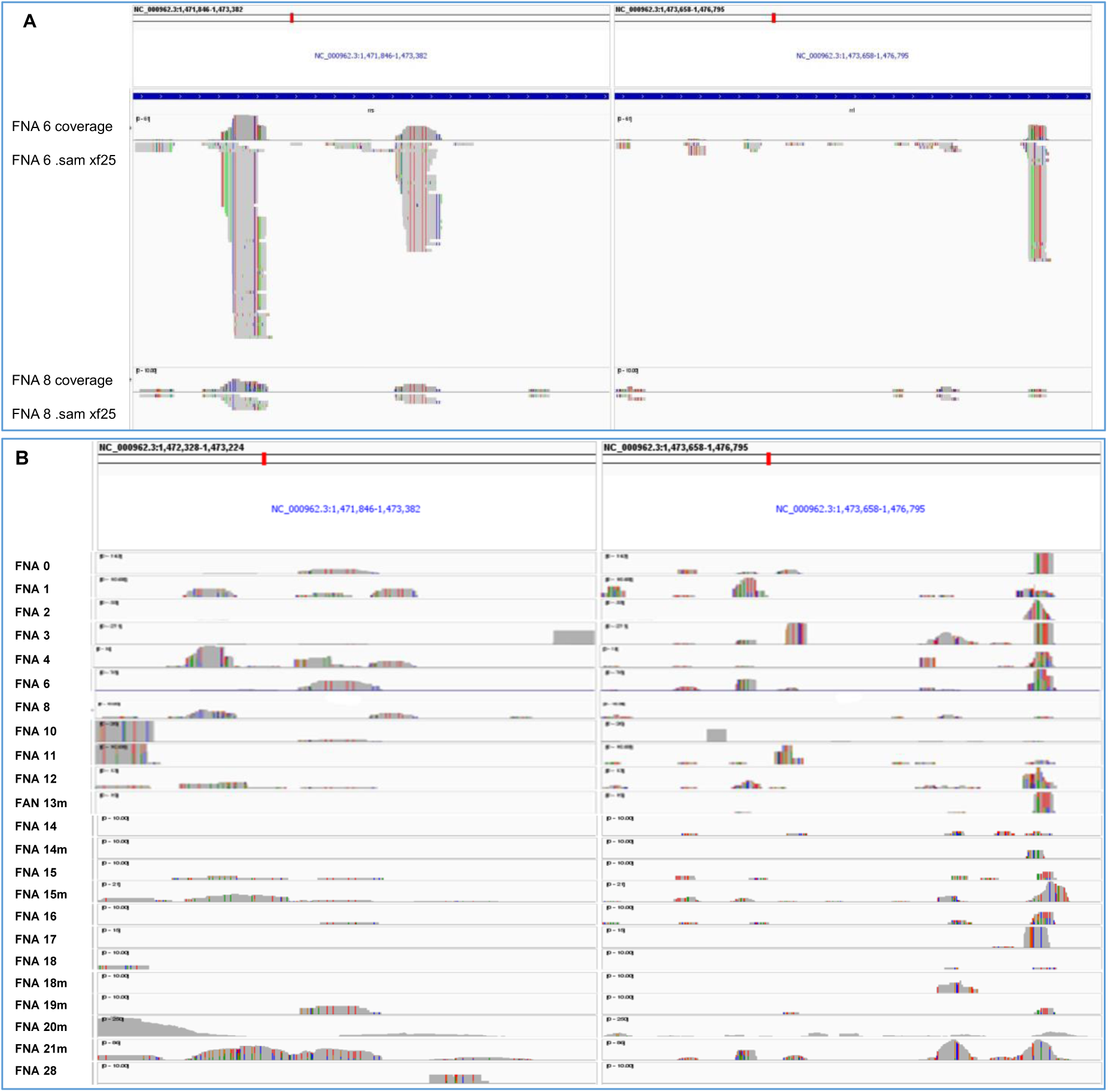
Coverage of rrs and rrl genes by Mtb reads detected in FNAs. No bacterial sequences were detected in FNA 22, 24, 25, 27. IGV alignment of Mtb reads associated with host cell UMIs from FNA 6 and FNA 8, respectively (**Panel A**), illustrates the variation between FNA 6 and FNA 8 in the numbers of transcript reads mapping to the small (rrs) and large (rrl) ribosomal RNA genes in the reference Mtb H37Rv genome. Multiple reads of identical length, starting position and directionality, usually in pairs or sets of three, are interpreted as arising from a single RNA source which we hypothesize was then amplified during library generation. **Panel B** shows the coverage of rrs and rrl genes in reads detected in the complete FNA set. Sequence variations versus the reference H37Rv genome are show in colors: G (brown), A (green), C (blue) and T (red).

**Fig. 10.**
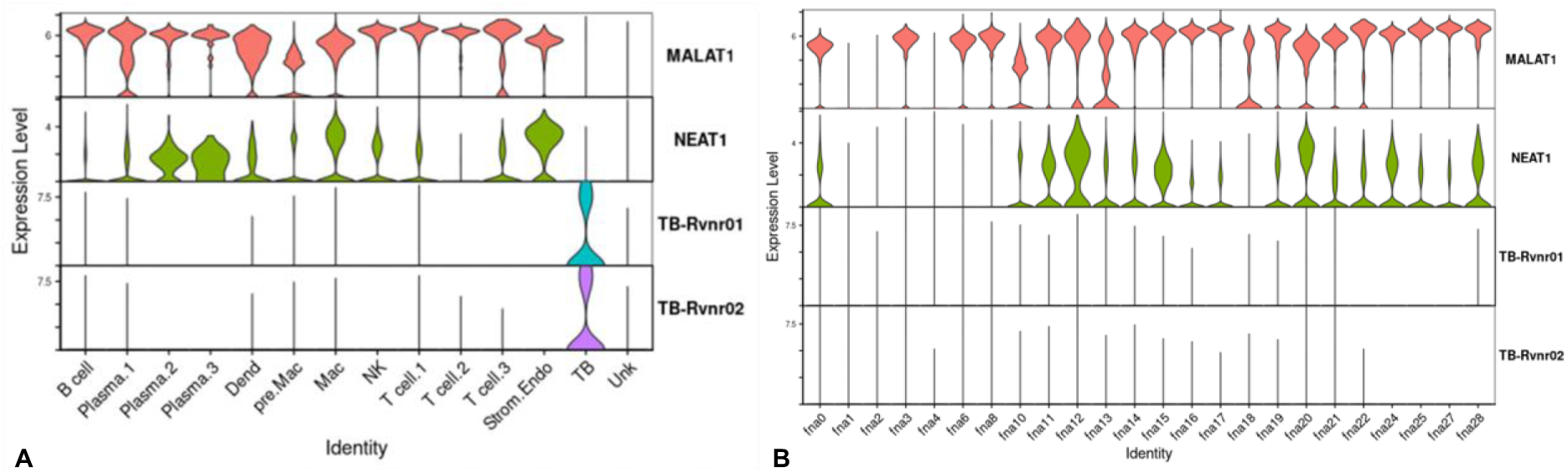
Expression of MALAT 1 in Combined Data Set. **Panel A** shows relative MALAT expression in the 14 clusters identified in figure 6. MALAT1 expression is significantly decreased in Mtb infected cells. **Panel B** shows expression of MALAT across all the analyzed FNAs. Only poor quality FNAs with small numbers of cells (e.g., FNA 1, 2 and 4) show low MALAT1 expression.

### Detection of HIV-1 infection in patient FNAs

Peripheral nodal tuberculosis granulomas appear in about 20% of patients, often in conjunction with pulmonary granulomas [33, 34]. The incidence is higher in women and in patients with HIV-1 co-infection [35-38]. Because HIV-1 is endemic to PNG at about 1% of the population, we included human HIV-1 genome (NC_001802.1) in the CellRanger genome assembly. Lymph nodes have been reported to be a site of HIV-1 persistence, even in undetectable patients [39-42]. In fact, we did detect rare HIV-1 transcripts in 3 of our 23 FNA samples (FNAs 18, 25 and 28), however, only the TAR sequence in the LTRs was detected; in one patient splicing into the nef gene (Fig. 11). TAR expression is often associated with HIV-1 latency. None of these 3 FNAs contained high levels of Mtb infected cells, as enumerated either by infected cell number or by the number of detected Mtb transcript reads.

**Fig. 11.**
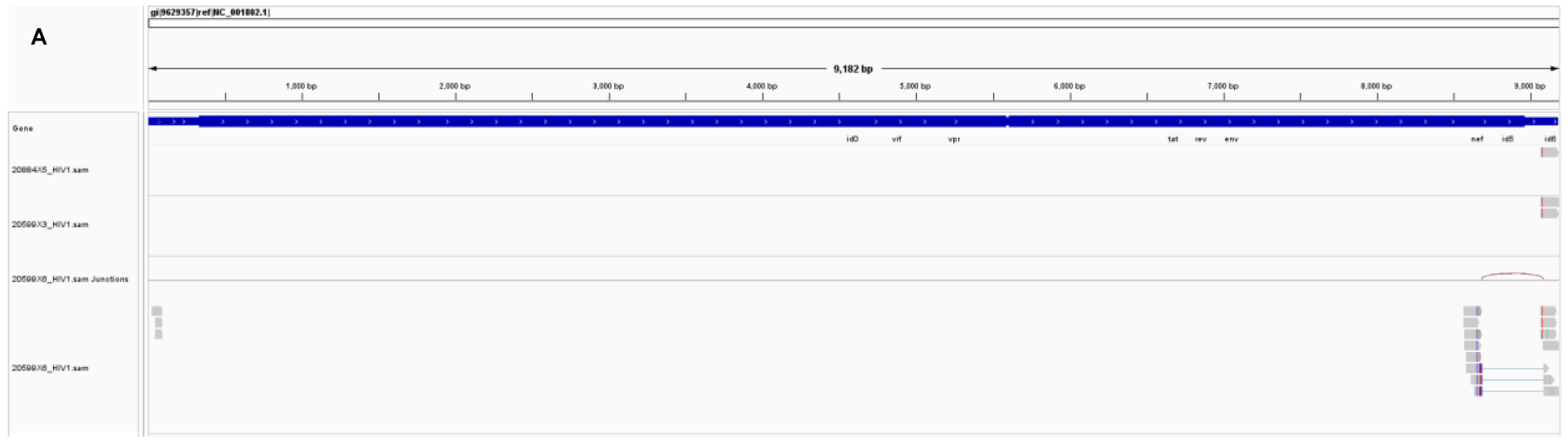
Integrated Genome Viewer Mapping of HIV-1 Transcript Reads Detected. **A)** HIV-1 reads detected in FNAs 18, 25 and 28 (from top to bottom).

## Discussion

We present here the first single cell RNA sequencing analysis of human fine needle aspirate samples from peripheral nodal lymphadenopathies associated with TB. We demonstrate such analysis is achievable in resource limited settings where higher incidences of tuberculosis often predominate. This proof-of-concept result provides new perspectives on the interactions between cells making up the presumed granulomatous lymphadenopathy. The data suggest that utilization of this innovative technology, with improvements in depth of library production and support of a state of the art microbiology laboratory, might soon provide important insights into patient status. Due to the work load at the tuberculosis clinic at POMGH, and the limited number of trained staff, there was no opportunity yet to obtain significant supporting pathology or microbiological follow up on the samples obtained here. There was confirmation of TB status of some of the FNAs by Pathology Laboratory microscopy of acid fast smears and/or GeneXpert analysis. Of our 23 de-identified FNA samples, 10 were confirmed as TB positive (FNAs 0, 4, 10, 13, 17, 18, 20, 21, 22 and 24). Multiple publications have reported ZN confirmation of TB status to be between 25% and 60% [19-21]. GeneXpert confirmation in similar resource limited settings is reported to be up to 70%, when available [19, 20]. None of the presenting patients were known to be HIV-1 positive.

We found that one pass of aspiration yielded ∼5 to 50 million cells for analysis in 1.25 mL 80% methanol, much in excess that is needed for scRNA seq analysis. With experience we improved our preservation protocol to include brief incubations with RBC lysis buffer and Accutase™ before methanol preservation, which yielded preserved FNA samples with suitable single cell suspensions after rehydration and the 10X Genomics scRNA seq pipeline. We look forward to new innovations in preservation for single cell analysis [43] which promise increased cell numbers and improved library quality and depth of coverage.

The cell types making up any one given FNA were assigned using our list of selective marker transcripts, in combination with SingleR [29], CellMarker_Augmented_2021 [30] and other [44] tools. Our list of selective gene transcripts was derived by laboriously vetting our own differential gene expression data, and from vetting many literature sources against our human samples. Many putative literature markers for cell identification, (e.g., for Th or M subtype cells) were not found to be sufficiently selective, or not sufficiently abundant in our data set to be of use in this regard. We included many more markers for monocyte/macrophage/dendrocyte type cells than the others because we found in our data set that often only limited numbers of the clustered putative macrophage/dendrocyte cells expressed our selective markers and often only one or two of the selective marker transcripts could be associated with that cluster. This was in contrast to the other clusters routinely identified in the individual UMAPS where CD3D and CD8A were found to be highly and selectively expressed in T cells, MS4A1 was found to be highly and selectively expressed in B cells, NKG7 was highly and selectively expressed in NK cells and JCHAIN and IGHG1 were highly and selectively expressed in plasma cells. These initial discriminating markers, in combination with the rest of the marker set gave confidence to cell type identification, which could then be tested for consistency with cell identifications generated with other tools. Further confirmation of the cell type identification (and quantification) was obtained later, when the cellular composition of the FNAs determined for the combined data set were shown to agree with that of the individual Loupe Browser analyses.

Further agreement was found when signaling data from the combined data set was correlated to the cells known to express those “Sender” or “Receiver” markers in the individual Loupe UMAPS. Thus, the individual Loupe UMAPS were consistent with the cell types known to express those signaling ligands or receptors. Cluster 0, a highly abundant cluster which was largely derived from FNAs 10, 18 and 20, and a few others, was assigned a pre-macrophage/dendrocyte identification because of low but consistent expression of several selective macrophage marker transcripts and because of numbers of immature macrophage lineage cells identified in that group by SingleR. This identification was also borne out by signaling interactions determined from CellChat.

Quantification of the given cell types comprising an individual FNA relied on the enumeration of cells in their respective clusters. We used K Means clustering tools to match Loupe Browser clusters to the cell types expressing genes on our feature list. These cluster populations were used to determine the percentage of total cells that cluster represented in a given FNA. On the occasions when K Means clustering divided or did not capture all the specifically identified cells of a given type, the Loupe Browser Lasso tool was used to capture the cells as a single cluster. For about half the FNAs (Fig. 2) the resolution T cells and NK cells was good enough for confidence in assigning separate clusters, although these clusters often abutted or transitioned from one to the other. We conducted repeated analyses which yielded consistent cluster population estimations, good enough for repetition of the trend analyses presented in figures 3 and 4.

However, many of the FNAs with small cell numbers or poorer library quality yielded intermingled clusters of T and NK cells, and occasionally intermingled B and plasma cells. In these cases, the intermingled clusters were designated T&NK or B&Pl (Fig. 1). In addition, when and individual cell type was particularly rare in a given FNA, such as in cases of FNAs with very few Mtb-infected cells, those cells often intermingled with various other named cell clusters and an estimation of their total count could only be obtained from the barcode count for marker being used (usually TB_sum). In FNAs with low Mtb counts, those Mtb-infected cells often associated with the plasma cell cluster, or less frequently with the macrophage/dendrocyte cluster. It was noted that, after Mtb-infected cells, plasma cells often had the lowest relative MALAT1 expression of the remaining cell types, which was perhaps indicative of other transcriptional similarities.

With confidence in cell type identification, we looked for trends or relationships between the cell types comprising the FNA samples (Fig. 3, 4). Extensive literature in human and NHP models suggests that cells making up the granuloma change with bacterial load and that this reflects mechanisms of bacterial control [22, 45, 46]. We initially looked for trends in cellular composition as we ranked FNAs according to increasing Mtb-infected cells, as determined by cluster analysis, or Mtb barcodes. We did not find any such significant trend in this limited data set. Nor did we identify any other significant trends until we focused on samples in which we could distinguish NK cell clusters from T cell clusters. From these higher quality samples, we noted an inverse relationship between the two cell types. Another, less significant trend was noted as well, the increase of macrophage/dendrocyte cells with increasing NK cell percentage (or decreasing T cell percentage).

To confirm the cell composition trends identified from Loupe Browser UMAP analyses (Fig. 3), cell cluster percentages determined by clustering in the combined data set was used. In this case, all the respective sub-clusters determined by the SEURAT clustering to be macrophage and dendrocyte (e.g., Clusters 0, 5 and 10), B (e.g., Clusters 3 and 9), T (e.g., Clusters 1, 7 and 8) or plasma lineage cells (e.g., Clusters 6, 11 and 14) were combined and used to determine cell type percentage composition of their respective source FNA. Thus, these FNA composition numbers were determined by the dimension reduction algorithm rather than the manual cluster identification process we used to determine FNA composition of individual Loupe Browser UMAPS. A bias introduced in this analysis is that FNAs with few cell number totals were given equal weight to FNAs with tens of thousands cells. Nevertheless, one significant trend confirmed for the total combined 18 FNAs was an inverse relationship between T cell percentages and macrophage/dendrocyte cells. The other previously noted trends were identifiable as well, but with less significance when using the full combined set of 18 FNAs. However, if the lower quality FNAs were excluded, the NK vs T and NK vs macrophage/dendrocyte cell trends were confirmed with strong confidence.

With the apparent confirmation of an inverse relationship between NK cells and T cells, we sought trends in our limited numbers of selective T cell marker transcripts. With increasing NK cell populations in a given FNA, it was found that T cells with CD8A or IL7R selective markers significantly proportionately decreased. This trend suggests a hypothesis where the pro-inflammatory role of T cells decreases with increasing NK cell participation (or vice versa). Future work with additional resources and orthogonal assessment techniques will be required to investigate this possibility.

Granulomatous tuberculosis lymph nodes are spatially organized structures, composed of a mixture of immune and non-immune cells [4, 5, 47]. However, this three dimensional structure is lost to our analysis. The fact that we routinely obtain the spectrum of cell types described as foundational to granuloma formation, and that we detect Mtb infected cells in those samples, argues that we are indeed sampling the essential granuloma structure, however without microscopy this assertion cannot be confirmed. We do not see several cell types reported in microscopic analysis of human or NHP studies in our FNAs [22, 23]. We do not see significant numbers of neutrophils. As postulated earlier, we believe this to be in part due to their low levels of transcription and lack of selective markers. In addition, Loupe Browser analysis of individual FNAs did not cluster high numbers of dendrocytes, as distinct from macrophages. While the analysis of differentially expressed transcripts in the combined data set clearly identified a dendrocyte cluster, these cells were few in number in most of the individual FNA UMAP analyses, and those few cells were most often clustered along with the more numerous macrophage/monocytic lineage cells. This yielded our terminology of macrophage/dendrocyte for the combined clusters. Endothelial and stromal cells, also identified in the combined data set, were only occasionally observed in the individual Loupe UMAPs, and never at high percentages. We hypothesize that the negative pressure on the needle during the FNA sampling biases for cells that are less adherent to the lymph node stroma or connective tissue. Thus, we are perhaps selectively vacuuming up the B, T, NK, plasma, macrophage/dendrocyte and Mtb-infected cells we identify here.

This pilot study shows a wide variation in the proportions of the cell types detected as making up the FNA samples. We cannot determine if the cellular variation we see in FNA composition is due to actual node to node differences amongst the patients, or if it is in part due to sampling differences amongst granulomas of varying structure. The majority of samples presented were obtained by our Registrar coauthors at UPNG (Masters of Medicine trainees in Pathology in the years 2019-2023). Future work and additional resources will be needed to answer this question more adequately. Furthermore, there was often significant contamination of the FNAs with erythrocytes, which seems inevitable considering the bloody nature of some of the aspirates. Once corrected for, the percentage of RBCs in a given FNA did not appear to affect the other correlations or analyses presented here, other than to reduce the total number of leukocytes analyzed. Removing the erythrocytic cells and cells with high MT gene expression from the combined data set did, however, significantly alter the cell-cell communications pathways identified by the CellChat analysis.

CellChat analysis was found to be insightful because it identified and ranked cell-cell communication ligands and receptors in the original combined data set. Many cytokines and chemokines have been identified as associated with Mtb infection [47, 48]. Interestingly, IL10, IL17 and INFγ, TNFα or their receptors, were not among those we identified. These are cytokines reported by previous foundational studies as pivotal to nodal granuloma function in humans and non-human primates [22,23]. A study by Kathmandu and colleagues [49] reported elevated baseline levels of IL-13 and IL-10, with reduced levels of IL-4 and GM-CSF in lymph nodes of Mtb infected individuals. They also reported increased levels of IFNγ, TNFα, IL-2, IL-17F, IL-22, IL-1α and GM-CSF, with reduced levels of TGFβ in primary lymph node cultures challenged with Mtb antigens. A report of Zhang and co-workers [50], using bioinformatics analysis of lymph node tuberculosis identified CXCL9, CD36 and LEP as genes involved in the pathogenesis of lymph node tuberculosis.

Instead, Collagen, MIF, MHC I, FN1, APP, Laminin, CD45, PECAM1&2 and CD99 were among the most highly expressed communication systems identified here. Many of these mediators play key roles in macrophage activation or maturation (MIF and APP), T cell activation and maturation (MHC I and CD45) and leukocyte migration (PECAM1 and 2). Many of the other identified ligands and receptors (Fig. 8) participate in cell-cell interactions related to adhesion and maturation (Collagen, FN1, Laminin, CD99, etc.). It was interesting to note that some of the most minor (by population number) sub-groups of dendrocytes, plasma cells and T cells were major contributors to the predominant signaling pathways identified by CellChat anlysis. Circular reasoning suggests that these signaling differences are why the sub-groups were clustered independently from the more populous clusters of those respective cell types in the first place. As mentioned before, when these signaling ligands or receptors were mapped in the individual Loupe UMAPS they all associated with their putative parent cell type, either “Sender” or “Receiver”. Almost all of these signaling interactions have immunomodulatory effects and are intuitively sensible, some having pro-inflammatory activities. We hypothesize that blocking or disruption of some of these key pathways could disrupt the structure or function of the granuloma and that the predominating signaling pathways in an individual’s FNA might provide insight as to that granuloma’s maturation or status.

We postulate that the detection of Mtb transcripts associated with host cell UMIs might provide a measure of Mtb burden in the FNA and presumably, the granuloma. Due to a lack of laboratory resources in Papua New Guinea, it has not yet been possible to see if the number of Mtb transcript reads, or if the number of detected infected cells, does indeed correlate with bacterial burden by colony forming unit (CFU) or chromosomal equivalent quantification (CEQ) analysis [51-52]. We hypothesize this will be the case. For all of our FNAs, the Mtb genome coverage of the detected bacterial transcript reads was insufficient to determine Mtb lineage. In the few cases where rrs or rrl coverage was extensive (e.g., FNAs 20 and 21), no sequence changes corresponding to potential drug resistance were identified when compared to the list in the “Catalogue of mutations in *Mycobacterium tuberculosis* complex and their association with drug resistance – Supplementary document” [53]. Single cell RNA sequencing of cell co-culture experiments with THP-1 and GFP-H37Ra suggest that we detect Mtb transcripts in less than 10% of the infected cells (our unpublished data). It is hoped that as scRNA seq precision and depth continue to improve that more complete coverage of the infecting Mtb genomes will be obtained. In future studies we plan to set aside a fraction of the FNA for CFU, CEQ analyses, and for liquid culture to enable WGS of the infecting pathogen(s).

We anticipate that there are likely complications of co-infection with other viral, bacterial or fungal pathogens in FNA samples such as those studied here. This possibility may be addressed using meta-genomic approaches following brief liquid culture of FNA aliquots. The complications introduced by HIV-1 co-infection might also be sorted out in studies that specifically include sufficient numbers of co-infected individuals. In our particular case, detection of HIV-1 co-infection was relayed back to the TB Clinic staff for follow up using their established protocols.

Overall, we believe this proof of concept study illuminates some of the potential insights that single cell analyses might provide into the structure and function of TB-associated lymphadenopathy. The continuing rapid improvements in these single cell technologies promise additional new understandings of host-pathogen interactions in the future.

## Materials and Methods

### FNA Sampling

Fine needle aspiration samples were collected as follows: after granting permission, suspected pLNTB patients with nodal enlargement ≥0.8 cm underwent 2 to 3 passes into the affected lymph nodes with a 22-gauge needle. Patient data is shown in figure S-7. No more than one lymph node per patient was sampled. A new needle was used for each pass. One aspirates was washed directly into 9.5 mL ice-cold RPMI buffer containing 0.2% fetal bovine serum, in a heparinized tube and mixed gently.

Our scRNA-seq preservation protocol required that the samples were processed as soon as possible after sampling. The ∼10 mL of the original FNA mixture was pelleted for 5 min. and resuspended on ice in 2 mL NH_4_Cl lysis solution [55] to lyse contaminating erythrocytes. After a maximum of 5 min. with occasional gentle mixing on ice, and observation of the depletion of obvious erythrocytes, 1 mL of Accutase™ was added directly to the lysis buffer for a maximum of an additional 3 min., again with occasional gentle mixing and observation of the dissolution of obvious tissue clots in the solution. The sample volume was expanded with 6 mL ice cold RPMI buffer and the cells pelleted again. The pelleted cells were gently suspended in 200 µl preservation buffer following 10X Genomics protocol [25] to which 1 mL of ice-cold methanol is slowly added with mixing. These preserved, sterilized, and de-identified samples were kept in the freezer or on ice packs for storage and transportation, followed by rehydration [25,57] and analysis by scRNA-seq.

### Single-cell RNA-sequencing

scRNA-seq was performed on single-cell suspensions using 10X Genomics Chromium to prepare cDNA sequencing libraries as described by Brady et al. [25, 57]. The samples were processed using the Chromium Single Cell 3′ V3 Kit (10X Genomics, Cat. # 1000075) using whole cells fixed in 80% methanol. Single cells were diluted to a target of 1000 cell/μL in 1× PBS (whole cells) or 1× PBS + 1.0% BSA + 0.2 U/μL RiboLock RNase Inhibitor to generate GEM’s prepared at a target of 10000 cells per sample. Barcoding, reverse transcription, and library preparation were performed according to manufacturer instructions. 10X Genomics generated cDNA libraries will be sequenced on NovaSeq 6000 instruments using 150 cycle paired-end sequencing at a depth of 10K reads per cell. The scRNA-seq was performed at the High Throughput Genomics Core at Huntsman Cancer Institute of the University of Utah.

For analytical procedures, the 10X Genomics Cell Ranger Single Cell software pipeline [25] is deployed to produce alignments and counts, utilizing the prescribed default parameters. The genomic references used for alignment were the human (hg38), the H37Rv Mtb (NC_00096.3:1) and HIV-1 (NC_001802.1). For quality management and further analytical exploration, Seurat (4.1.0) was utilized. Doublets were identified with DoubletFinder, cells were excluded based on having less than 100 genes/features and an excess of 25% mitochondrial genes. Mitochondrial genes were filtered out but every cell that contained Mtb genes was retained. Dimensionality was reduced and scaled via SCTransformation (0.3.5) using the Gamma-Poisson generalized linear model (glmGamPoi, 1.4.0) methodology at default resolution or less. Automated categorization of cells was performed using SingleR (1.6.1). Statistics within Seurat pipelines were generated with FindAllMarkers or FindMarkers which utilizes a Wilcoxon rank sum test [30; 57-68].

## Acknowledgements

The authors wish to recognize the funding supporting for this work: the “Comprehensive mapping of TB/HIV-1 nodal granuloma in humans” Skaggs Foundation Research Grant, and the University of Utah Seed Grant: “Understanding HIV persistence one cell at a time”. AFC recognizes support from K08 AI139339 and the University of Utah Department of Pathology start-up funds. The authors also wish to acknowledge Professor Nakapi Tefuarani, Dean UPNG School of Medicine for his enduring support, and Dr. Rodney Itaki and Dr. Evelyn Lavu (decd), UPNG School of Medicine for early assistance in obtaining our IRB. We also wish to acknowledge the excellent support obtained through the University of Utah Core facilities: "Research reported in this publication utilized the High-Throughput Genomics and Cancer Bioinformatics Shared Resource at Huntsman Cancer Institute at the University of Utah and was supported by the National Cancer Institute of the National Institutes of Health under Award Number P30CA042014. The content is solely the responsibility of the authors and does not necessarily represent the official views of the NIH."

## Notes

### Competing Interest Statement

The authors have declared no competing interest.

### Summary of Updates

We found errors in the original reference list, which has been updated and corrected here.

## References

1 Ruby Maini; Shivaraj Nagalli. Stat Pearls[Internet] Lymphadenopathy https://www.ncbi.nlm.nih.gov/books/NBK558918/

2 Asano S. Granulomatous lymphadenitis. J Clin Exp Hematop. 2012;52(1):1-16. doi: 10.3960/jslrt.52.1. PMID: 22706525.

3 Lyon SM, Rossman MD. Pulmonary tuberculosis. Microbiol Spectr. (2017) 5:1–13. doi: 10.1128/microbiolspec.TNMI7-0032-2016

4 Granulomatous Diseases of the Head and Neck. Medscape. [cited 2024 Jan 30] Available from: https://emedicine.medscape.com/article/854739-overview?form=fpf.

5 Granuloma. Cleveland Clinic. [cited 2024 Jan 30] Available from: https://my.clevelandclinic.org/health/diseases/24597-granuloma.

6 Latent TB Infection and TB Disease. [cited 2024 Jan 30] Available from: https://www.cdc.gov/tb/topic/basics/tbinfectiondisease.htm.

7 Ehlers, S., and Schaible, U.E. (2013). The granuloma in tuberculosis: dynamics of a host-pathogen collusion. Front. Immunol. 3, 411. 10.3389/fimmu.2012.00411.

8 Guirado E, Schlesinger L. Modeling the Mycobacterium tuberculosis granuloma–the critical battlefield in host immunity and disease. Front Immunol. (2013) 4:1–7. doi: 10.3389/fimmu.2013.00098

9 Shah KK, Pritt BS, Alexander MP. Histopathologic review of granulomatous inflammation. J Clin Tuberc Other Mycobact Dis. 2017 Feb 10;7:1–12. doi: 10.1016/j.jctube.2017.02.001. PMID: 31723695; PMCID: PMC6850266.

10 Pagan, A.J., and Ramakrishnan, L. (2014). Immunity and immunopathology in the tuberculous granuloma. Cold Spring Harb. Perspect. Med. 5, a018499. 10.1101/cshperspect.a018499.

11 Ganchua SKC, White AG, Klein EC, Flynn JL (2020) Lymph nodes—The neglected battlefield in tuberculosis. PLoS Pathog 16(8): e1008632. 10.1371/journal.ppat.1008632Lymph nodes are a principle cite of bacterial control infection

12 Silva Miranda M, Breiman A, Allain S, Deknuydt F, Altare F. The tuberculous granuloma: an unsuccessful host defence mechanism providing a safety shelter for the bacteria? Clin Dev Immunol. 2012;2012:139127. doi: 10.1155/2012/139127. Epub 2012 Jul 3. PMID: 22811737; PMCID: PMC3395138.

13. WHO. Global tuberculosis report 2021. [cited 2024 Jan 30]. Available from: https://www.who.int/publications/i/item/9789240037021

14 Daniel TM. The history of tuberculosis. Respir Med. 2006; 100(11):1862–70. Epub 2006/09/05. 10.1016/j.rmed.2006.08.006 PMID: 16949809.

15. Latent TB Infection and TB Disease. [cited 2024 Jan 30]. Available from: https://www.cdc.gov/tb/topic/basics/tbinfectiondisease.htm

16 Houben RM, Dodd PJ. The Global Burden of Latent Tuberculosis Infection: A Re-estimation Using Mathematical Modelling. PLoS Med. 2016; 13(10):e1002152. Epub 2016/10/26. 10.1371/journal.pmed. 1002152 PMID: 27780211; PubMed Central PMCID: PMC5079585.

17 Itaki R, Bannick, Lavu E, Joseph J, Magaye R, Banamu J, Johnson K, Welch H. 2016. Drug susceptibility pattern of Mycobacterium tuberculosis isolates from patients undergoing fine needle aspiration biopsy at Port Moresby General Hospital. Pacific Journal of Medical Sciences 15: 26–33.

18. Itaki RL. 2016. MMED Dissertation Diagnostic accuracy of Xpert ® MTB/RIF using AFB microscopy and cytomorphology as reference tests in the diagnosis of tuberculous lymphadenitis inpatients presenting for fine needle aspiration biopsy at the Port Moresby General Hospital.

19 Negi SS, Khan SF, Gupta S, Pasha ST, Khare S, Lal S. Comparison of the conventional diagnostic modalities, bactec culture and polymerase chain reaction test for diagnosis of tuberculosis. Indian J Med Microbiol. 2005 Jan;23(1):29–33. doi: 10.4103/0255-0857.13869. PMID: 15928418

20 Ngabonziza JC, Ssengooba W, Mutua F, Torrea G, Dushime A, Gasana M, Andre E, Uwamungu S, Nyaruhirira AU, Mwaengo D, Muvunyi CM. Diagnostic performance of smear microscopy and incremental yield of Xpert in detection of pulmonary tuberculosis in Rwanda. BMC Infect Dis. 2016 Nov 8;16(1):660. doi: 10.1186/s12879-016-2009-x. PMID: 27825314; PMCID: PMC5101805.

21 Dzodanu EG, Afrifa J, Acheampong DO, Dadzie I. Diagnostic Yield of Fluorescence and Ziehl-Neelsen Staining Techniques in the Diagnosis of Pulmonary Tuberculosis: A Comparative Study in a District Health Facility. Tuberc Res Treat. 2019 Apr 10;2019:4091937. doi: 10.1155/2019/4091937. PMID: 31093372; PMCID: PMC6481031.

22 Gideon HP, Hughes TK, Tzouanas CN, Wadsworth MH 2nd, Tu AA, Gierahn TM, Peters JM, Hopkins FF, Wei JR, Kummerlowe C, Grant NL, Nargan K, Phuah JY, Borish HJ, Maiello P, White AG, Winchell CG, Nyquist SK, Ganchua SKC, Myers A, Patel KV, Ameel CL, Cochran CT, Ibrahim S, Tomko JA, Frye LJ, Rosenberg JM, Shih A, Chao M, Klein E, Scanga CA, Ordovas-Montanes J, Berger B, Mattila JT, Madansein R, Love JC, Lin PL, Leslie A, Behar SM, Bryson B, Flynn JL, Fortune SM, Shalek AK. Multimodal profiling of lung granulomas in macaques reveals cellular correlates of tuberculosis control. Immunity. 2022 May 10;55(5):827–846.e10. doi: 10.1016/j.immuni.2022.04.004. Epub 2022 Apr 27. PMID: 35483355; PMCID: PMC9122264.

23 Diedrich CR, O’Hern J, Gutierrez MG, Allie N, Papier P, Meintjes G, Coussens AK, Wainwright H, Wilkinson RJ. Relationship Between HIV Coinfection, Interleukin 10 Production, and Mycobacterium tuberculosis in Human Lymph Node Granulomas. J Infect Dis. 2016 Nov 1;214(9):1309–1318. doi: 10.1093/infdis/jiw313. Epub 2016 Jul 26. PMID: 27462092; PMCID: PMC5079364.

24. 24 Sholeye AR, Williams AA, Loots DT, Tutu van Furth AM, van der Kuip M and Mason S (2022) Tuberculous Granuloma: Emerging Insights From Proteomics and Metabolomics. Front. Neurol. 13:804838. doi: 10.3389/fneur.2022.804838

25. 10X Genomics Available from: https://www.10xgenomics.com/?userresearcharea=ra_g&utm_medium=search&gclid=Cj0KCQjwoeemBhCfARIsADR2QCsCe34XQhIvVaogRsruoLYmbTA7IAk5Jrp3D8fQRwzUEudsj8MsrHMaAgqbEALw_wcB&usercampaignid=7011P000001XhgOQAS&userregion=multi&useroffertype=website-page&utm_source=google&utm_campaign=sem-goog-2023-05-website-page-brand-2023-brand-sem-programs-paid-search-7890

26 Zhong Y, Ashley CL, Steain M, Ataide SF. Assessing the suitability of long non-coding RNAs as therapeutic targets and biomarkers in SARS-CoV-2 infection. Front Mol Biosci. 2022 Aug 16;9:975322. doi: 10.3389/fmolb.2022.975322. PMID: 36052163; PMCID: PMC9424846.

27 Huang K, Wang C, Vagts C, Raguveer V, Finn PW, Perkins DL. Long non-coding RNAs (lncRNAs) NEAT1 and MALAT1 are differentially expressed in severe COVID-19 patients: An integrated single cell analysis. medRxiv [Preprint]. 2021 Jul 31:2021.03.26.21254445. doi: 10.1101/2021.03.26.21254445. Update in: PLoS One. 2022 Jan 10;17(1):e0261242. PMID: 33821282; PMCID: PMC8020982.

28 The Human Protein Atlas. Available from: https://www.proteinatlas.org/

29 Hu C, Li T, Xu Y, Zhang X, Li F, Bai J, Chen J, Jiang W, Yang K, Ou Q, Li X, Wang P, Zhang Y. CellMarker 2.0: an updated database of manually curated cell markers in human/mouse and web tools based on scRNA-seq data. Nucleic Acids Res. 2023 Jan 6;51(D1):D870–D876. doi: 10.1093/nar/gkac947. PMID: 36300619; PMCID: PMC9825416.

30 Bioconductor SingleR Available from: doi:10.18129/B9.bioc.SingleR

31 Jin S, Plikus MV, Nie Q. CellChat for systematic analysis of cell-cell communication from single-cell and spatially resolved transcriptomics bioRxiv 2023.11.05.565674; doi: 10.1101/2023.11.05.565674

32 Huang K, Wang C, Vagts C, Raguveer V, Finn PW, Perkins DL. Long non-coding RNAs (lncRNAs) NEAT1 and MALAT1 are differentially expressed in severe COVID-19 patients: An integrated single-cell analysis. PLoS One. 2022 Jan 10;17(1):e0261242. doi: 10.1371/journal.pone.0261242. PMID: 35007307; PMCID: PMC8746747.

33 Alami NN, Yuen CM, Miramontes R, Pratt R, Price SF, Navin TR; Centers for Disease Control and Prevention (CDC). Trends in tuberculosis - United States, 2013. MMWR Morb Mortal Wkly Rep. 2014 Mar 21;63(11):229–33. PMID: 24647398; PMCID: PMC4584631.

34 Rieder HL, Snider DE Jr, Cauthen GM. Extrapulmonary tuberculosis in the United States. Am Rev Respir Dis. 1990 Feb;141(2):347–51. doi: 10.1164/ajrccm/141.2.347. PMID: 2301852.

35 Naing C, Mak JW, Maung M, Wong SF, Kassim AI. 2013 Meta-analysis: the association between HIV infection and extrapulmonary tuberculosis. Lung. 191:27–34. doi: 10.1007/s00408-012-9440-6. Epub 2012 Nov

36 Shivakoti R, Sharma D, Mamoon G, Pham K 2017 Association of HIV infection with extrapulmonary tuberculosis: a systematic review Infection.45:11–21. doi:10.1007/s15010-016-0960-5.

37 Rolo M, González-Blanco B, Reyes CA, Rosillo N, López-Roa P. Epidemiology and factors associated with Extra-pulmonary tuberculosis in a Low-prevalence area. J Clin Tuberc Other Mycobact Dis. 2023 May 12;32:100377. doi: 10.1016/j.jctube.2023.100377. PMID: 37252369; PMCID: PMC10209530.

38 Min J, Park JS, Kim HW, Ko Y, Oh JY, Jeong YJ, Na JO, Kwon SJ, Choe KH, Lee WY, Lee SS, Kim JS, Koo HK. Differential effects of sex on tuberculosis location and severity across the lifespan. Sci Rep. 2023 Apr 13;13(1):6023. doi: 10.1038/s41598-023-33245-5. PMID: 37055508; PMCID: PMC10102118.

39 Banga R, Munoz O, Perreau M. HIV persistence in lymph nodes. Curr Opin HIV AIDS. 2021 Jul 1;16(4):209–214. doi: 10.1097/COH.0000000000000686. PMID: 34059608.

40 McManus WR, Bale MJ, Spindler J, Wiegand A, Musick A, Patro SC, Sobolewski MD, Musick VK, Anderson EM, Cyktor JC, Halvas EK, Shao W, Wells D, Wu X, Keele BF, Milush JM, Hoh R, Mellors JW, Hughes SH, Deeks SG, Coffin JM, Kearney MF. HIV-1 in lymph nodes is maintained by cellular proliferation during antiretroviral therapy. J Clin Invest. 2019 Jul 30;129(11):4629–4642. doi: 10.1172/JCI126714. PMID: 31361603; PMCID: PMC6819093.

41 Yukl SA, Kaiser P, Kim P, Telwatte S, Joshi SK, Vu M, Lampiris H, Wong JK. HIV latency in isolated patient CD4^+^ T cells may be due to blocks in HIV transcriptional elongation, completion, and splicing. Sci Transl Med. 2018 Feb 28;10(430):eaap9927. doi: 10.1126/scitranslmed.aap9927. PMID: 29491188; PMCID: PMC5959841.

42 Rosen EP, Deleage C, White N, Sykes C, Brands C, Adamson L, Luciw P, Estes JD, Kashuba ADM. Antiretroviral drug exposure in lymph nodes is heterogeneous and drug dependent. J Int AIDS Soc. 2022 Apr;25(4):e25895. doi: 10.1002/jia2.25895. PMID: 35441468; PMCID: PMC9018350.

43 10x Genomics Support/Single Cell Gene Expression Flex/Documentation/Sample Prep/Fixation of Cells & Nuclei for Chromium Fixed RNA Profiling. https://www.10xgenomics.com/support/single-cell-gene-expression-flex/documentation/steps/sample-prep/fixation-of-cells-and-nuclei-for-chromium-single-cell-gene-expression-flex. Jan. 2024

44 Ianevski A, Giri AK, Aittokallio T. Fully-automated and ultra-fast cell-type identification using specific marker combinations from single-cell transcriptomic data. Nat Commun. 2022 Mar 10;13(1):1246. doi: 10.1038/s41467-022-28803-w. PMID: 35273156; PMCID: PMC8913782.

45 McCaffrey EF, Donato M, Keren L, Chen Z, Delmastro A, Fitzpatrick MB, Gupta S, Greenwald NF, Baranski A, Graf W, Kumar R, Bosse M, Fullaway CC, Ramdial PK, Forgó E, Jojic V, Van Valen D, Mehra S, Khader SA, Bendall SC, van de Rijn M, Kalman D, Kaushal D, Hunter RL, Banaei N, Steyn AJC, Khatri P, Angelo M. The immunoregulatory landscape of human tuberculosis granulomas. Nat Immunol. 2022 Feb;23(2):318–329. doi: 10.1038/s41590-021-01121-x. Epub 2022 Jan 20. Erratum in: Nat Immunol. 2022 May;23(5):814. PMID: 35058616; PMCID: PMC8810384.

46 Lin PL, Ford CB, Coleman MT, Myers AJ, Gawande R, Ioerger T, Sacchettini J, Fortune SM, Flynn JL. Sterilization of granulomas is common in active and latent tuberculosis despite within-host variability in bacterial killing. Nat Med. 2014 Jan;20(1):75–9. doi: 10.1038/nm.3412. Epub 2013 Dec 15. PMID: 24336248; PMCID: PMC3947310.

47 Marakalala MJ, Raju RM, Sharma K, Zhang YJ, Eugenin EA, Prideaux B, Daudelin IB, Chen PY, Booty MG, Kim JH, Eum SY, Via LE, Behar SM, Barry CE 3rd, Mann M, Dartois V, Rubin EJ. Inflammatory signaling in human tuberculosis granulomas is spatially organized. Nat Med. 2016 May;22(5):531–8. doi: 10.1038/nm.4073. Epub 2016 Apr 4. PMID: 27043495; PMCID: PMC4860068.

48 Domingo-Gonzalez R, Prince O, Cooper A, Khader SA. Cytokines and Chemokines in Mycobacterium tuberculosis Infection. Microbiol Spectr. 2016 Oct;4(5):10.1128/microbiolspec.TBTB2-0018-2016. doi: 10.1128/microbiolspec.TBTB2-0018-2016. PMID: 27763255; PMCID: PMC5205539.

49 Kathamuthu GR, Moideen K, Sridhar R, Baskaran D, Babu S. Enhanced Mycobacterial Antigen-Induced Pro-Inflammatory Cytokine Production in Lymph Node Tuberculosis. Am J Trop Med Hyg. 2019 Jun;100(6):1401–1406. doi: 10.4269/ajtmh.18-0834. PMID: 30994092; PMCID: PMC6553923.

50 Zhang Z, Liu Y, Wang W, Xing Y, Jiang N, Zhang H, Zhang H, He L, Yue W, Jiang L, Wang K. Identification of Differentially Expressed Genes Associated with Lymph Node Tuberculosis by the Bioinformatic Analysis Based on a Microarray. J Comput Biol. 2020 Jan;27(1):121–130. doi: 10.1089/cmb.2019.0161. Epub 2019 Aug 28. PMID: 31460784.

51 Muñoz-Elías EJ, Timm J, Botha T, Chan WT, Gomez JE, McKinney JD. Replication dynamics of Mycobacterium tuberculosis in chronically infected mice. Infect Immun. 2005 Jan;73(1):546–51. doi: 10.1128/IAI.73.1.546-551.2005. PMID: 15618194; PMCID: PMC538940.

52 Lin, P.L., Coleman, T., Carney, J.P.J., Lopresti, B.J., Tomko, J., Fillmore, D., Dartois, V., Scanga, C., Frye, L.J., Janssen, C., et al. (2013). Radiologic responses in cynomolgus macaques for assessing tuberculosis chemotherapy regimens. Antimicrob. Agents Chemother. 57, 4237– 4244. 10.1128/AAC.00277-13.

53 WHO Catalogue of mutations in Mycobacterium tuberculosis complex and their association with drug resistance – Supplementary document. WHO/UCN/GTB/PCI/2021.7-©WHO 2021. CC BY-NC-SA 3.0 IGO licence. https://www.who.int/publications/i/item/9789240028173

54 Cadena AM, Hopkins FF, Maiello P, Carey AF, Wong EA, Martin CJ, Gideon HP, DiFazio RM, Andersen P, Lin PL, Fortune SM, Flynn JL. Concurrent infection with Mycobacterium tuberculosis confers robust protection against secondary infection in macaques. PLoS Pathog. 2018 Oct 12;14(10):e1007305. doi: 10.1371/journal.ppat.1007305. PMID: 30312351; PMCID: PMC6200282.

55 University of Washington RBC Lysing Solutions and Cell Lysing Procedure https://depts.washington.edu/flowlab/Cell%20Analysis%20Facility/RBC%20Lysing%20Solutions%20and%20Cell%20Lysing%20Procedure.pdf

56 Brady SW, McQuerry JA, Qiao Y, Piccolo SR, Shrestha G, Jenkins DF, Layer RM, Pedersen BS, Miller RH, Esch A, Selitsky SR, Parker JS, Anderson LA, Dalley BK, Factor RE, Reddy CB, Boltax JP, Li DY, Moos PJ, Gray JW, Heiser LM, Buys SS, Cohen AL, Johnson WE, Quinlan AR, Marth G, Werner TL, Bild AH. Combating subclonal evolution of resistant cancer phenotypes. Nat Commun. 2017 Nov 1;8(1):1231. doi: 10.1038/s41467-017-01174-3.

57 Hafemeister C, Satija R. Normalization and variance stabilization of single-cell RNA-seq data using regularized negative binomial regression. Genome Biol. 2019 Dec 23;20(1):296. doi: 10.1186/s13059-019-1874-1. PMID: 31870423; PMC6927181.

58 hlmann-Eltze C, Huber W. glmGamPoi: fitting Gamma-Poisson generalized linear models on single cell count data. Bioinformatics. 2021 Apr 5;36(24):5701–5702. doi: 10.1093/bioinformatics/btaa1009. PMID: 33295604;

59 Hao Y, Hao S, Andersen-Nissen E, Mauck WM 3rd, Zheng S, Butler A, Lee MJ, Wilk AJ, Darby C, Zager M, Hoffman P, Stoeckius M, Papalexi E, Mimitou EP, Jain J, Srivastava A, Stuart T, Fleming LM, Yeung B, Rogers AJ, McElrath JM, Blish CA, Gottardo R, Smibert P, Satija R. Integrated analysis of multimodal single-cell data. Cell. 2021 Jun 24;184(13):3573–3587.e29. doi: 10.1016/j.cell.2021.04.048. Epub 2021 May 31. PMID: 34062119; PMCID: PMC8238499.

60 Aran D, Looney AP, Liu L, Wu E, Fong V, Hsu A, Chak S, Naikawadi RP, Wolters PJ, Abate AR, Butte AJ, Bhattacharya M. Reference-based analysis of lung single-cell sequencing reveals a transitional profibrotic macrophage. Nat Immunol. 2019 Feb;20(2):163–172. doi: 10.1038/s41590-018-0276-y. Epub 2019 Jan 14. PMID: 30643263; PMCID: PMC6340744.

61 Stuart T, Butler A, Hoffman P, Hafemeister C, Papalexi E, Mauck WM 3rd, Hao Y, Stoeckius M, Smibert P, Satija R. Comprehensive Integration of Single-Cell Data. Cell. 2019 Jun 13;177(7):1888–1902.e21. doi: 10.1016/j.cell.2019.05.031. Epub 2019 Jun 6. PMID: 31178118; PMCID: PMC6687398.

62. Bodenhofer U, Kothmeier A, Hochreiter S. APCluster: an R package for affinity propagation clustering. Bioinformatics. 2011 Sep 1;27(17):2463–4. doi: 10.1093/bioinformatics/btr406. Epub 2011 Jul 6. PMID: 21737437.

63 Liao Y, Smyth GK, Shi W. The Subread aligner: fast, accurate and scalable read mapping by seed-and-vote. Nucleic Acids Res. 2013 May 1;41(10):e108. doi: 10.1093/nar/gkt214. Epub 2013 Apr 4. PMID: 23558742; PMCID: PMC3664803.

64 Shen Y, Rahman M, Piccolo SR, Gusenleitner D, El-Chaar NN, Cheng L, Monti S, Bild AH, Johnson WE. ASSIGN: context-specific genomic profiling of multiple heterogeneous biological pathways. Bioinformatics. 2015 Jun 1;31(11):1745–53. doi: 10.1093/bioinformatics/btv031. Epub 2015 Jan 22. PMID: 25617415; PMCID: PMC4443674.

65 Home Bioconductor 3.17 Software Packages scran Available from: https://bioconductor.org/packages/release/bioc/html/scran.html

66 Lun ATL, Riesenfeld S, Andrews T, Dao TP, Gomes T; participants in the 1st Human Cell Atlas Jamboree; Marioni JC. EmptyDrops: distinguishing cells from empty droplets in droplet-based single-cell RNA sequencing data. Genome Biol. 2019 Mar 22;20(1):63. doi: 10.1186/s13059-019-1662-y. PMID: 30902100; PMCID: PMC6431044.

67 Srinivasula SM, Ashwell JD. IAPs: what’s in a name? Mol Cell. 2008 Apr 25;30(2):123–35. doi: 10.1016/j.molcel.2008.03.008. PMID: 18439892; PMCID: PMC2677451.

68 loess: Local Polynomial Regression Fitting. Available from: Accessed 02/02/2022. https://rdrr.io/r/stats/loess.html

