## Supplementary Information for "Single Cell Analysis of Peripheral TB-Associated Granulomatous Lymphadenitis"

### Supplemental Figures

| Clusters | FNAs | | | | | | | | | | | | | | | | | | | | | | | |
|---|---|---|---|---|---|---|---|---|---|---|---|---|---|---|---|---|---|---|---|---|---|---|---|---|
| | fna 0 | fna 1 | fna 10 | fna 11 | fna 12 | fna 13 | fna 14 | fna 15 | fna 16 | fna 17 | fna 18 | fna 19 | fna 2 | fna 20 | fna 21 | fna 22 | fna 24 | fna 25 | fna 27 | fna 28 | fna 3 | fna 4 | fna 6 | fna 8 |
| T Cell | | | | 584 | | | 2734 | 6346 | | 6838 | | 724 | | | | 421 | 1754 | 1894 | 13398 | 412 | | | | |
| B Cell | 471 | | | 193 | 7 | 102 | 1936 | 463 | 1681 | 2780 | 711 | 240 | | | 1367 | 154 | 509 | 1591 | 7573 | 149 | | | | |
| Macrophage/ Dendrocyte | 159 | | 645 | 59 | 123 | 191 | 1210 | 1267 | 57 | 341 | 4840 | 536 | | 2579 | 47 | 86 | 437 | 46 | 102 | 248 | | | 7 | |
| Plasma Cell | 62 | | | 66 | 2 | 68 | 200 | 2879 | 207 | 413 | 637 | 17 | | 303 | 110 | 271 | 93 | 21 | 1804 | 19 | | | | |
| NK Cell | | | | 420 | | | 2103 | 2741 | | 789 | | 2070 | | | | 199 | 449 | 488 | 3560 | 610 | | | | |
| TB sum | 114 | 23 | 21 | 152 | 50 | 103 | 15 | 33 | 12 | 20 | 3 | 0 | 34 | 696 | 229 | 1 | 0 | 0 | 0 | 1 | 67 | 28 | 152 | 16 |
| T cell/NK | 859 | | 465 | 1004 | 57 | 717 | 4841 | 9087 | 5314 | | 1966 | | | 303 | 674 | | | | | | | | 1295 | |
| B cell/Plasma cell | | | 50 | | | | 2163 | 3342 | | | | | | 305 | 1477 | | | | | | | | 1258 | |
| | | | | | | 1746 UNK/ | 15329 RBC | | | | 2032 RBC | | | | | | | | | | | | | |
| Total Bar Codes Sum with Max unless empty | 1808 | 3925 | 1118 | 1466 | 231 | 2855 | 23626 | 13593 | 6929 | 11418 | 7695 | 5656 | 3925 | 3609 | 2212 | 1126 | 3173 | 4075 | 26804 | 1426 | 3708 | 14883 | 2710 | 13678 |

**S-1** Sum of 1st and max by K means clustering of total cell counts to match distinguishing features with Lasso tool and barcodes when necessary, from Fig. 2.

All granulomas combined

Res 0.1 201, 828 cells passed QC – all TB and HIV cells preserved

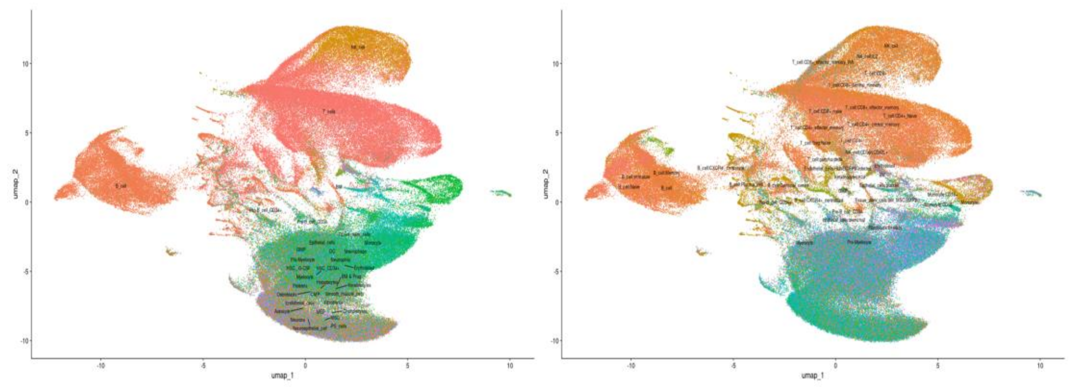


S-2 **Left)** Combined Data Set Resolution 0.1, SingleR Main. **Right)** Combined Data Set Resolution 0.1, SingleR Fine.

| Clusters | FNAs | | | | | | | | | | | | | | | | | | | | | | | |
|---|---|---|---|---|---|---|---|---|---|---|---|---|---|---|---|---|---|---|---|---|---|---|---|---|
| | fna 0 | fna 1 | fna 10 | fna 11 | fna 12 | fna 13 | fna 14 | fna 15 | fna 16 | fna 17 | fna 18 | fna 19 | fna 2 | fna 20 | fna 21 | fna 22 | fna 24 | fna 25 | fna 27 | fna 28 | fna 3 | fna 4 | fna 6 | fna 8 |
| 0 pre-M | 74 | 29 | 64570 | 5 | 2 | 100 | 10 | 44 | 6 | 14 | 2302 | 12 | 150 | 1455 | 27 | 2 | 13 | 0 | 188 | 0 | 7 | 566 | 11 | 2 |
| 1 T1 | 843 | 0 | 716 | 665 | 32 | 486 | 2846 | 7053 | 2461 | 6927 | 1593 | 2139 | 25 | 202 | 701 | 509 | 1718 | 547 | 14897 | 375 | 861 | 38 | 1008 | 749 |
| 2 Unk | 0 | 0 | 0 | 0 | 0 | 0 | 0 | 0 | 0 | 0 | 0 | 0 | 0 | 0 | 0 | 0 | 0 | 0 | 0 | 0 | 0 | 0 | 0 | 0 |
| 3 B | 447 | 0 | 98 | 179 | 8 | 70 | 1793 | 434 | 1480 | 2750 | 636 | 685 | 2 | 4 | 1317 | 143 | 503 | 1534 | 6646 | 134 | 540 | 2 | 1047 | 72 |
| 4 NK | 29 | 0 | 78 | 297 | 8 | 96 | 1624 | 808 | 30 | 89 | 134 | 7353 | 0 | 33 | 17 | 66 | 188 | 102 | 384 | 570 | 8 | 2 | 20 | 24 |
| 5 M | 36 | 0 | 687 | 74 | 137 | 214 | 976 | 922 | 47 | 141 | 34 | 2754 | 8 | 1854 | 32 | 105 | 487 | 25 | 219 | 252 | 6 | 4 | 8 | 43 |
| 6 Pl1 | 136 | 1 | 1169 | 65 | 1 | 45 | 92 | 2206 | 113 | 590 | 865 | 21 | 0 | 4 | 75 | 194 | 75 | 37 | 2961 | 0 | 6 | 2 | 36 | 6 |
| 7 T2 | 2 | 0 | 8 | 2 | 12 | 0 | 0 | 24 | 2452 | 0 | 0 | 8 | 0 | 0 | 2 | 16 | 0 | 23 | 309 | 0 | 0 | 0 | 0 | 0 |
| 8 T3 | 4 | 59 | 408 | 0 | 0 | 27 | 0 | 3 | 2 | 6 | 33 | 1 | 63 | 4 | 1 | 0 | 1 | 1610 | 9 | 0 | 0 | 94 | 1 | 1 |
| 9 D | 83 | 0 | 154 | 14 | 6 | 39 | 30 | 805 | 50 | 421 | 3 | 28 | 1 | 26 | 48 | 13 | 56 | 27 | 87 | 3 | 0 | 1 | 7 | 0 |
| 10 S&E | 36 | 0 | 8 | 0 | 6 | 0 | 1 | 362 | 9 | 15 | 1 | 68 | 0 | 7 | 2 | 8 | 1 | 0 | 13 | 1 | 0 | 0 | 6 | 0 |
| 11 Pl2 | 0 | 0 | 0 | 0 | 0 | 0 | 0 | 351 | 0 | 0 | 0 | 0 | 0 | 0 | 0 | 0 | 0 | 0 | 0 | 0 | 0 | 0 | 0 | 0 |
| 12 TB | 8 | 3 | 10 | 2 | 7 | 3 | 4 | 0 | 0 | 0 | 0 | 2 | 1 | 80 | 11 | 0 | 0 | 0 | 0 | 1 | 41 | 4 | 59 | 7 |
| 13 Pl3 | 0 | 0 | 0 | 0 | 0 | 0 | 0 | 34 | 0 | 0 | 0 | 0 | 0 | 0 | 0 | 0 | 0 | 0 | 0 | 0 | 0 | 0 | 0 | 0 |
| sum | 1698 | 92 | 67906 | 1303 | 219 | 1080 | 7376 | 13046 | 6650 | 10953 | 5601 | 13071 | 250 | 3669 | 2233 | 1056 | 3042 | 3905 | 25713 | 1336 | 1469 | 713 | 2203 | 904 |

S-3 Total cell counts for 14 clusters in Combined Data Set shown in Fig. 6.

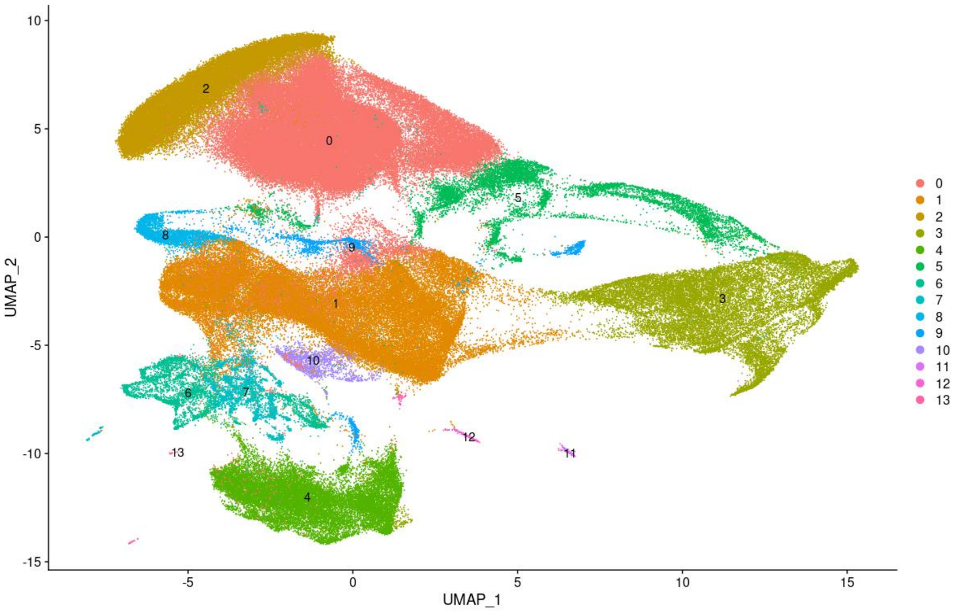


S-4 Original Combined Data Set Res. 0.1 used for CellChat v1.

7

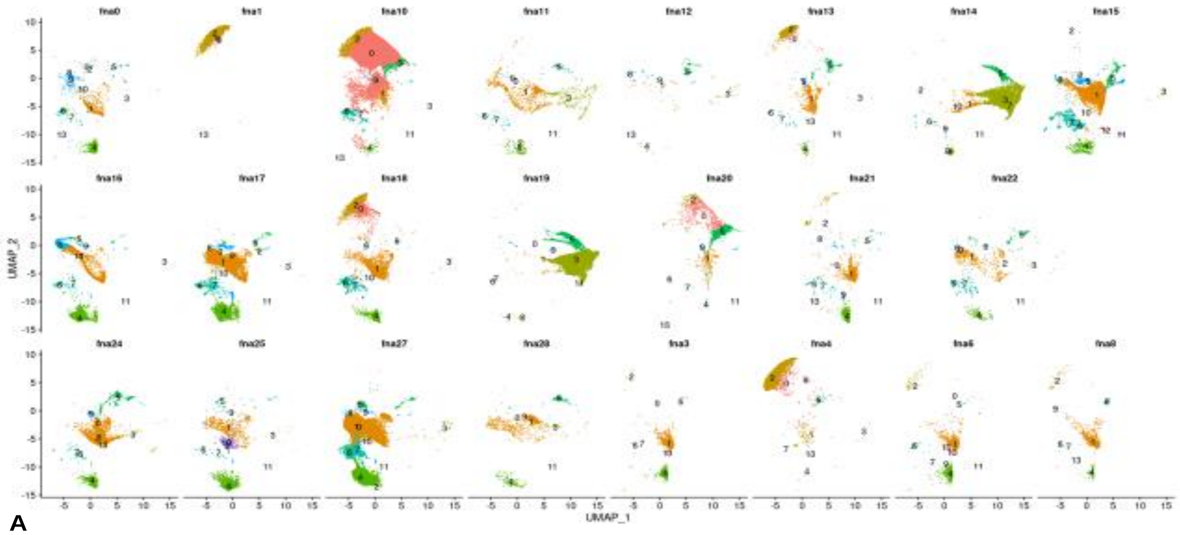


A

| Clusters | FNAs | | | | | | | | | | | | | | | | | | | | | | | |
|---|---|---|---|---|---|---|---|---|---|---|---|---|---|---|---|---|---|---|---|---|---|---|---|---|
| | fna 0 | fna 1 | fna 10 | fna 11 | fna 12 | fna 13 | fna 14 | fna 15 | fna 16 | fna 17 | fna 18 | fna 19 | fna 2 | fna 20 | fna 21 | fna 22 | fna 24 | fna 25 | fna 27 | fna 28 | fna 3 | fna 4 | fna 6 | fna 8 |
| 0 pre-M | 74 | 29 | 64570 | 5 | 2 | 100 | 10 | 44 | 6 | 14 | 2302 | 12 | 150 | 1455 | 27 | 2 | 13 | 0 | 188 | 0 | 7 | 566 | 11 | 2 |
| 1 T1 | 843 | 0 | 716 | 665 | 32 | 486 | 2846 | 7053 | 2461 | 6927 | 1593 | 2139 | 25 | 202 | 701 | 509 | 1718 | 547 | 14897 | 375 | 861 | 38 | 1008 | 749 |
| 2 Unk | 0 | 0 | 0 | 0 | 0 | 0 | 0 | 0 | 0 | 0 | 0 | 0 | 0 | 0 | 0 | 0 | 0 | 0 | 0 | 0 | 0 | 0 | 0 | 0 |
| 3 B | 447 | 0 | 98 | 179 | 8 | 70 | 1793 | 434 | 1480 | 2750 | 636 | 685 | 2 | 4 | 1317 | 143 | 503 | 1534 | 6646 | 134 | 540 | 2 | 1047 | 72 |
| 4 NK | 29 | 0 | 78 | 297 | 8 | 96 | 1624 | 808 | 30 | 89 | 134 | 7353 | 0 | 33 | 17 | 66 | 188 | 102 | 384 | 570 | 8 | 2 | 20 | 24 |
| 5 M | 36 | 0 | 687 | 74 | 137 | 214 | 976 | 922 | 47 | 141 | 34 | 2754 | 8 | 1854 | 32 | 105 | 487 | 25 | 219 | 252 | 6 | 4 | 8 | 43 |
| 6 Pl1 | 136 | 1 | 1169 | 65 | 1 | 45 | 92 | 2206 | 113 | 590 | 865 | 21 | 0 | 4 | 75 | 194 | 75 | 37 | 2961 | 0 | 6 | 2 | 36 | 6 |
| 7 T2 | 2 | 0 | 8 | 2 | 12 | 0 | 0 | 24 | 2452 | 0 | 0 | 8 | 0 | 0 | 2 | 16 | 0 | 23 | 309 | 0 | 0 | 0 | 0 | 0 |
| 8 T3 | 4 | 59 | 408 | 0 | 0 | 27 | 0 | 3 | 2 | 6 | 33 | 1 | 63 | 4 | 1 | 0 | 1 | 1610 | 9 | 0 | 0 | 94 | 1 | 1 |
| 9 D | 83 | 0 | 154 | 14 | 6 | 39 | 30 | 805 | 50 | 421 | 3 | 28 | 1 | 26 | 48 | 13 | 56 | 27 | 87 | 3 | 0 | 1 | 7 | 0 |
| 10 S&E | 36 | 0 | 8 | 0 | 6 | 0 | 1 | 362 | 9 | 15 | 1 | 68 | 0 | 7 | 2 | 8 | 1 | 0 | 13 | 1 | 0 | 0 | 6 | 0 |
| 11 Pl2 | 0 | 0 | 0 | 0 | 0 | 0 | 0 | 351 | 0 | 0 | 0 | 0 | 0 | 0 | 0 | 0 | 0 | 0 | 0 | 0 | 0 | 0 | 0 | 0 |
| 12 TB | 8 | 3 | 10 | 2 | 7 | 3 | 4 | 0 | 0 | 0 | 0 | 2 | 1 | 80 | 11 | 0 | 0 | 0 | 0 | 1 | 41 | 4 | 59 | 7 |
| 13 Pl3 | 0 | 0 | 0 | 0 | 0 | 0 | 0 | 34 | 0 | 0 | 0 | 0 | 0 | 0 | 0 | 0 | 0 | 0 | 0 | 0 | 0 | 0 | 0 | 0 |
| sum | 1698 | 92 | 67906 | 1303 | 219 | 1080 | 7376 | 13046 | 6650 | 10953 | 5601 | 13071 | 250 | 3669 | 2233 | 1056 | 3042 | 3905 | 25713 | 1336 | 1469 | 713 | 2203 | 904 |

B

S-5 Original Combined Data Set Res. 0.1 used for CellChat v1.

S-6 CellChat v1 of Original Combined Data Set

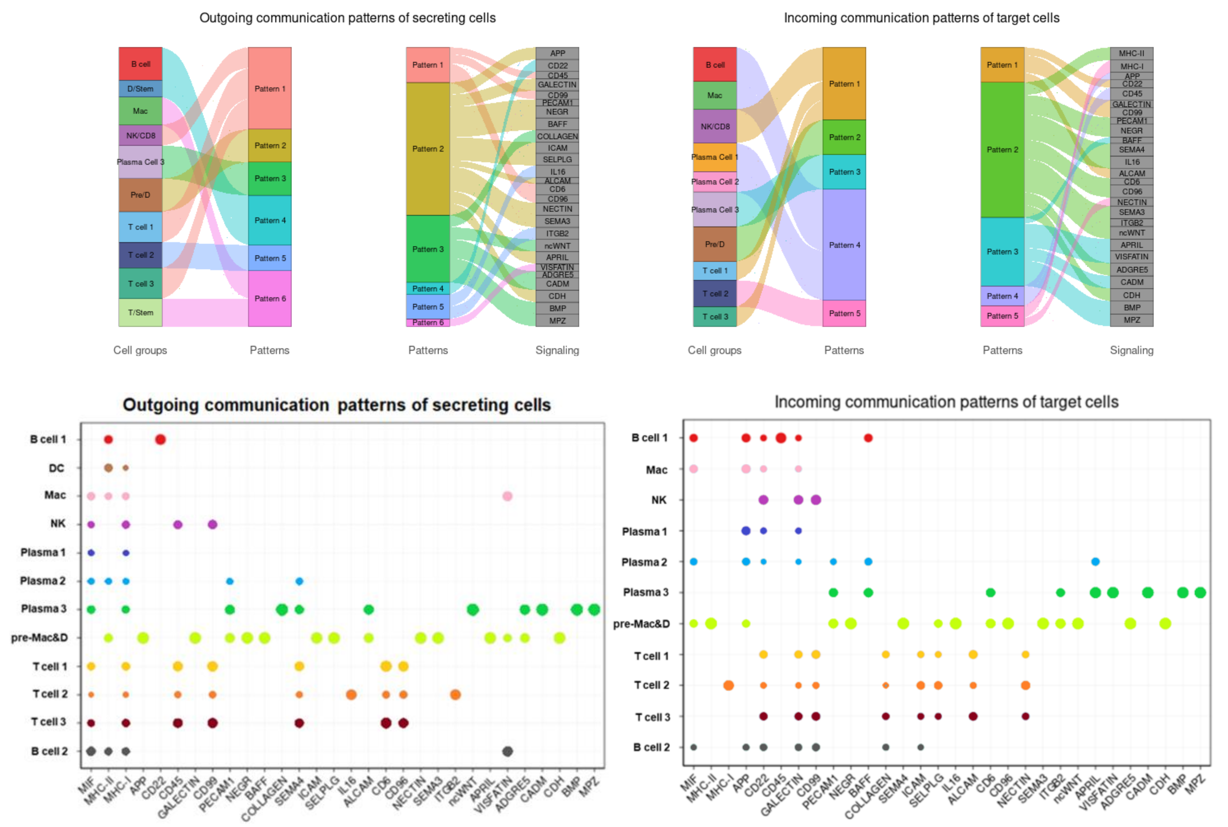

FNA Path Reports

| Sample | Sex | Age | TB+ Confirmed | Year |
|---|---|---|---|---|
| FNA0 | M | 22 | Y | 2019 |
| FNA1 | M | 36 | ND | 2022 |
| FNA2 | M | 38 | ND | 2022 |
| FNA3 | M | 41 | ND | 2022 |
| FNA4 | F | 27 | Y | 2022 |
| FNA6 | M | 17 | Y | 2022 |
| FNA8 | M | 3 | N | 2022 |
| FNA10 | M | 14 | Y | 2023 |
| FNA11 | F | 17 | N | 2023 |
| FNA12 | F | 23 | N | 2023 |
| FNA13 | M | 23 | Y | 2023 |
| FNA14 | F | 27 | N | 2023 |
| FNA15 | F | 35 | N | 2023 |
| FNA16 | F | 18 | N | 2023 |
| FNA17 | F | 20 | N/RC | 2023 |
| FNA18 | F | 35 | Y | 2023 |
| FNA19 | M | 56 | N | 2023 |
| FNA20 | F | 35 | Y/RC | 2023 |
| FNA21 | F | 28 | Y/RC | 2023 |
| FNA22 | F | 14 | Y | 2023 |
| FNA24 | M | 61 | Y | 2023 |
| FNA25 | M | 36 | N | 2023 |
| FNA27 | M | 41 | ND | 2023 |
| FNA28 | M | 69 | ND | 2023 |

Y=Yes; N=No; ND=Not Determined; RC=Recurrent

S-Table 1 Patient/Sample data; TB confirmation by POMGH TB Clinic Pathology Lab using ZN staining and/or GeneXpert molecular analysis.
